# Nitrite reductase NirB mediates an unconventional nitrogen assimilation strategy to enhance adaptations of extremophile

**DOI:** 10.64898/2026.09.23.753953

**Authors:** Ruirui Liu, Ye Tian

**Affiliations:** State Key Laboratory of Biocontrol, School of Life Sciences, Sun Yat-Sen University, Guangzhou, China

**Keywords:** nitrite reductase, dual NIR, nitrogen assimilation, adaptation

## Abstract

Microbial metabolism in extreme environments has long been shrouded in profound mystery. Distinct from the direct assimilation of exogenous organic carbon, microorganisms predominantly exploit inorganic nitrogen substrates as their primary nitrogen sources. As a critical nitrogen assimilation pathway, assimilatory nitrite reduction plays a vital role in microbial nitrogen acquisition, especially in extreme environments with poor nutrient availability. In this study, we identified the coexistence of two isoforms of NADH-dependent nitrite reductase in single genomes. NIR-α and NIR-β, named based on their phylogenetic relationships, exhibit significant differences in protein sequence, genomic context and evolutionary relationship, a phenomenon that has rarely been reported before. They also show distinct expression patterns, which may allow bacteria to achieve efficient nitrogen conversion in nutrient-poor environments. This finding calls for a revision of the conventional model of nitrogen cycling gene combinations inferred from common model microorganisms and holds numerous implications for how we understand and study the environmental adaptation mechanisms of extremophiles.

## Introduction

The Earth’s biosphere is dominated by cold environments, which cover approximately 80% of the planet’s surface, although this differs from our intuitive perception^1,2^. Cold environments extend far beyond the well-known Arctic and Antarctic regions, including approximately 70% of the global oceans, as well as high-altitude areas and deep-sea hadal zones^3–7^. Microorganisms that thrive in these cold environments are usually classified as psychrophiles^8–10^. As the most widely distributed extremophiles, they play a crucial role in biogeochemical cycling, which is critical for maintaining planetary fertility and habitability^11–17^. From a long-term perspective, studying microbial communities and their material-energy metabolism in cold biospheres is of great significance for understanding the vitality of Earth.

The *Psychromonas* group can serve as an indicator taxon for evaluating energy input and availability in cold aquatic environments, reflecting the strength of energy flow into the community^18^. As a typical genus in cold habitats, its widespread distribution encompasses various cold environments, including the polar regions, sea ice, and even deep-sea hadal habitats^19–24^. The distinctive membrane lipid composition and specialized proteins and metabolic pathways enable them to thrive in extreme conditions^25–28^. Notably, *Psychromonas* exhibits robust carbohydrate metabolic capacity, playing a pivotal role in carbon cycling within cold aquatic environments^29–32^. An in situ enrichment study in the Arctic Ocean revealed that *Psychromonas* is a key chitin-degrading group and can serve as an indicator taxon for chitinous input^33^. Furthermore, *Psychromonas* can utilize a range of algal polysaccharides, including sialic acid, mannitol, and fucoidan, which serve as significant organic reservoirs and are central to marine carbon cycling^34–38^. Taken together, *Psychromonas* appears to function primarily as a decomposer of marine organic carbon, with its significant role largely confined to the marine carbon cycle^30^.

As a matter of fact, *Psychromonas* may play multiple ecological roles beyond common perception. Recent research on marine wastewater treatment provided insight: *Psychromonas* accounted for the highest relative abundance (45.6%) in marine activated sludge within sequencing batch reactors, indicating its critical involvement and high efficiency in biological nitrogen removal from marine wastewater^39^. More direct evidence came from the identification of functional genes related to nitrogen cycling, which further supports the expanded ecological role of *Psychromonas*. For instance, a *nosZ-*specific fragment was amplified from the *P. ingrahamii* strain using specific primers, suggesting that it has the potential for denitrification^40, 41^. Another single-cell sequencing analysis of hadal samples confirmed that *Psychromonas* possesses the *nap* gene cluster (*napBDEF*), which mediates the reduction of nitrate to nitrite with the core protein NapA^42^. In addition, the potential for nitrogen fixation is demonstrated by the presence of a nitrogenase (*nif*) operon, identified in a *Psychromonas* strain isolated from the gut microbiome of isopods in Puerto Rico Trench^43^. Moreover, evidence indicated that in seawater with high abundance of *Psychromonas*, not only was organic carbon degradation markedly enhanced, but nitrogen cycling processes were also highly active^44^. In conclusion, these findings suggest that *Psychromonas* may play a significant role in nitrogen cycling, which seems particularly significant in oceanic systems because nitrogen content and availability constrain marine productivity^45^. However, a comprehensive understanding of the primary pathways and metabolic characteristics of *Psychromonas* in the nitrogen cycle remains limited, potentially underestimating its importance in cold environments.

To address the role of *Psychromonas* in the nitrogen cycle, we performed a comparative genomic analysis of the genus. The result revealed its gene repertoire, genomic plasticity and evolutionary history. We found that nitrogen metabolism in *Psychromonas* is primarily centered on nitrate reduction and nitrite-to-ammonium conversion, with a minority of genomes also encoding the potential for nitrogen fixation or denitrification. Unexpectedly, we identified the coexistence of two evolutionarily distinct NADH-dependent nitrite reductase systems (NIR systems) within the same genome, a dual isoform configuration that has not been reported to our knowledge. Subsequently, we characterized the sequence features, phylogenetic relationships and genomic contexts of both isoforms, and used protein structure modeling to identify key residues involved in ligand recognition in the large subunit, thereby providing insights into the catalytic mechanism. Moreover, expression analyses under different conditions further revealed the potential evolutionary plasticity and suggested that the dual NIR feature facilitates nitrogen accumulation under cold and nutrient-poor conditions, thereby enhancing environmental adaptability. In summary, our study identified a rare isoform combination in *Psychromonas* nitrogen cycling and provided new insight into the mechanisms by which psychrophiles adapt to environments.

## Results

### Phylogenetic analysis and genomic landscape of *Psychromonas*

For this study, we utilized *Psychromonas* genomes from GenBank, all of which met the quality standards of >95% completeness and <5% contamination, as outlined in **Supplementary Table 1**. The genus *Psychromonas* has an average genome size of 4.43 Mbp (from 3.98 to 5.53 Mbp) and a G+C content of 36.5-42.4%, with 3,201-3,776 protein-coding genes (**Figure 1a**). Notably, *P. aquimarina* and *P. ossibalaenae* have genomes larger than 5 Mbp and the highest G+C content among the taxa analyzed. To accurately resolve the phylogeny of *Psychromonas*, we constructed a maximum-likelihood tree based on a concatenated alignment of 120 marker genes, using *Colwellia* as the outgroup. The resulting phylogenomic tree showed that *Psychromonas* is monophyletic with high bootstrap support.

**Figure 1.**
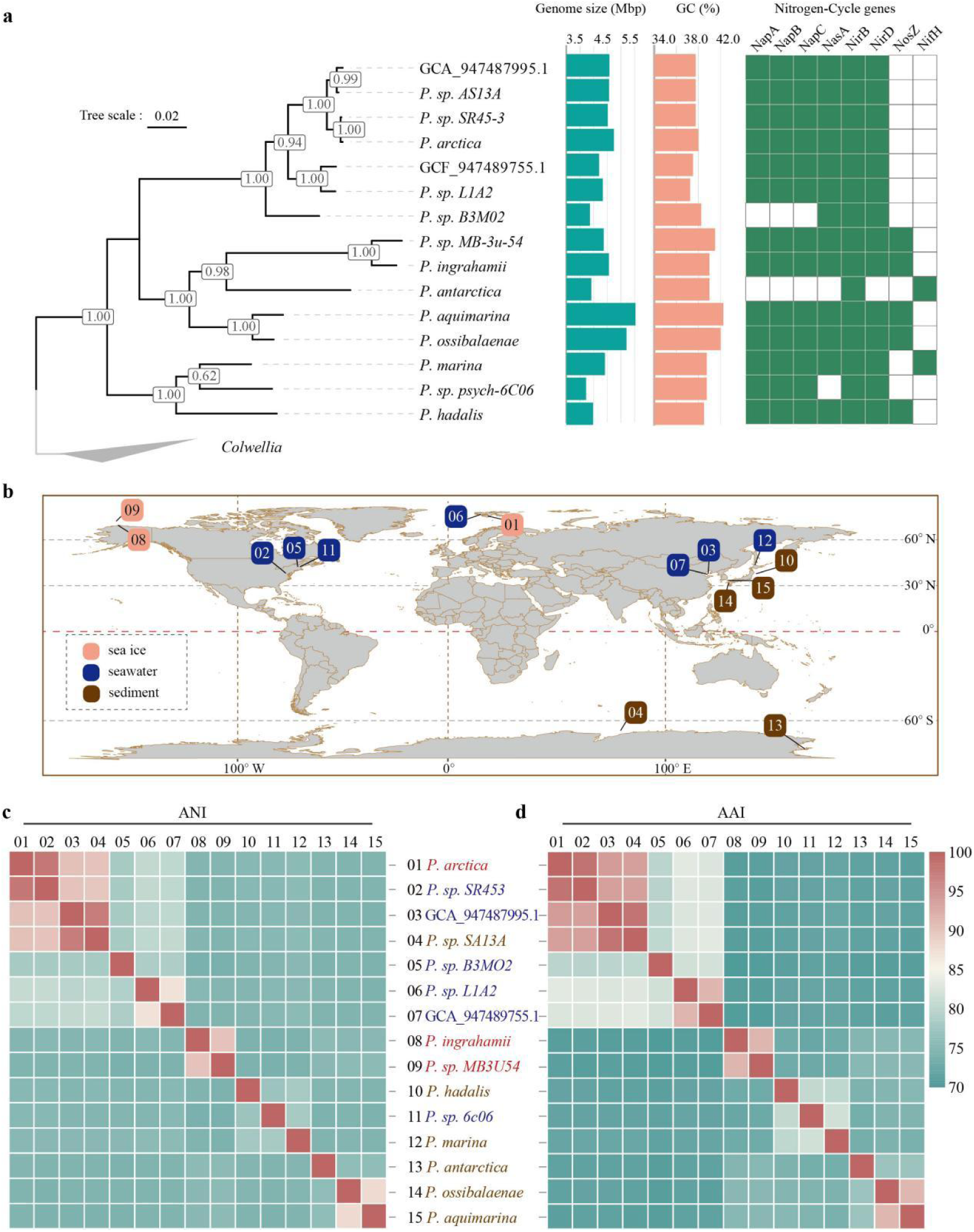
Phylogeny, distribution and genomic features of the *Psychromonas*. **(a)** Concatenated marker gene phylogeny of the genus *Psychromonas*. *Colwellia* was used as the outgroup and is shown as a grey clade. Bootstrap values are shown in square boxes at each node. Tree annotations indicate, from left to right, genome size (Mbp), GC content (%) and the presence (green squares) or absence (blank squares) of N-cycle pathways. **(b)** Geographic distribution of *Psychromonas* genomic data. Boxes indicate the sampling locations of the selected genomes. Colors represent different sample types. **(c)** Average nucleotide identity (ANI) and average amino acid identity (AAI) among *Psychromonas* species. Corresponding values are provided in Supplementary Tables 3. Numbers in **b** and **c** refer to the same set of genomes.

The geographic distribution of the collected genomes showed a clear pattern of concentration in mid- to high-latitude regions (**Figure 1b**); specifically, the collection included 60% of the samples from mid-latitudes (30-60°) and 40% from high latitudes (>60°), primarily from the Antarctic and Arctic. All samples were associated with cold habitats (**Supplementary Fig. 1 and Supplementary Table 2**). Several genomes were derived from deep-sea environments, including *Psychromonas hadalis*, which was isolated from trench sediment collected at a depth of 7,542 m^46^. It is reasonable to propose that these extreme conditions have driven the evolution of corresponding adaptive mechanisms in *Psychromonas*. Whole-genome average nucleotide identity (ANI) and average amino acid identity (AAI) analyses were also performed to evaluate genetic discontinuity and divergence among *Psychromonas* genomes (**Figure 1c and d; also see Supplementary Table 3**). Both metrics reflect shared ancestry and provide a natural framework for grouping genomes at the species and genus levels^47–49^. The high ANI and AAI values (>95%) confirmed that *Psychromonas* sp. SR453 and *P. arctica* constitute the same species, a conclusion consistent with their close phylogenetic relationship. Similarly, *Psychromonas* sp. SA13A and GCA_947487995.1 clustered into another minimal taxonomic unit, with the latter derived from an uncultured environmental sample. In addition, three other genomes pairs—GCA_947489755.1/*Psychromonas* sp. L1A2, *P. ingrahamii*/*Psychromonas* sp. MB3U54, and *P. ossibalaenae*/*P. aquimarina*—each exhibited distinct clustering, yet their ANI/AAI values were below 95%. The remaining genomes exhibited substantial differences in composition, with an average ANI of approximately 78% and corresponding AAI values ranging from 70% to 80%. These results indicate that the analyzed genomes represent at least thirteen *Psychromonas* species, providing a comprehensive landscape of genomic diversity within the genus.

On this basis, we screened *Psychromonas* genomes for nitrogen cycle-related genes to elucidate their potential roles in this process. All *Psychromonas* genomes contained genes associated with nitrogen metabolism (**Figure 1a and Supplementary Fig. 2**). Periplasmic dissimilatory nitrate reductase (Nap) and cytoplasmic assimilatory nitrate reductase (Nas) were detected in approximately 87% of genomes and are responsible for reducing nitrate to nitrite. The NADH-dependent nitrite reductase large subunit NirB and small subunit NirD were detected in 100% and 93% of genomes, respectively. They catalyze the reduction of nitrite to ammonium, which is subsequently incorporated into organic matter, thereby linking inorganic and organic nitrogen pools. Nitrous oxide reductase (NosZ), the only known biological sink responsible for N_2_O consumption, was present in approximately 33% of *Psychromonas* genomes. Additionally, the core nitrogenase component NifH was found restricted to *P. antarctica* and *P. marina*, although they are located on non-adjacent branches of the phylogenetic tree. In summary, these results indicate that the main nitrogen-utilization pathway in *Psychromonas* involves nitrate reduction to nitrite and subsequent assimilation into ammonium. Importantly, nitrite reductase NirB was present in all genomes, including compact genomes such as *Psychromonas* sp. psych-6C06 and *Psychromonas* sp. B3M02, highlighting its indispensable role in nitrogen cycling in *Psychromonas*.

### Nitrogen-cycling genes as a key component of the *Psychromonas* core genome

To obtain more comprehensive evidence linking nitrogen cycling genes to *Psychromonas* adaptation, we established a pan-genome analysis pipeline to interrogate the evolutionary signatures left by strong selective pressures in extreme environments. For this purpose, we reconstructed the *Psychromonas* pan-genome and identified 1,361 core orthogroups among 13,804 gene clusters (**Figure 2a; also see Supplementary Table 4**). Subsequently, to assess how the pan-genome composition changes with the inclusion of additional genomes, we computed rarefaction curves and performed data modelling based on Power Law and Heaps’ Law^50^. The fitted Power Law alpha value was 0.599 ± 0.0013, indicating that additional orthogroups would continue to be identified as more genomes are sampled. This inference was also supported by the Heaps’ Law gamma parameter (0.484 ± 0.003), and both lines of evidence suggest that the *Psychromonas* pan-genome is open. Moreover, the core-genome curve showed clear convergence as strain sampling increased, indicating that our dataset sufficiently captured the conserved genetic repertoire of the genus (**Figure 2a**). It is the core genome that captures the most fundamental survival requirements of *Psychromonas*, serving as the backbone that supports the remainder of the genome^51^. To better understand the functions of the *Psychromonas* genome, the eggNOG database was used for orthology analysis to achieve functional annotation of *Psychromonas* genomes (see Methods)^52^. The properties and statistics of genes assigned to clusters of orthologous groups (COG) functional categories are shown in **Figure 2b and Supplementary Table 5**. The most enriched categories were E (amino acid transport and metabolism) and P (inorganic ion transport and metabolism), respectively. These categories include genes for nitrogen uptake and metabolism—processes critical for survival in cold environments—and ultimately reflect the shaping force of natural selection on the genome. Similar gene counts were found for energy production and conversion (C) and carbohydrate transport and metabolism (G), a pattern consistent with the chemoheterotrophic lifestyle of *Psychromonas*. Furthermore, the numbers of genes assigned to categories K and L were also strongly correlated, possibly resulting from the existence of a transcription-replication interaction profile (TRIP), whereby DNA replication can shape genome-wide gene expression^53^. A lower number of genes was commonly observed in the remaining functional categories (e.g., D and V). Clearly, pathways related to nitrogen metabolism were the most significantly enriched, highlighting their predominant representation within the *Psychromonas* genome.

**Figure 2.**
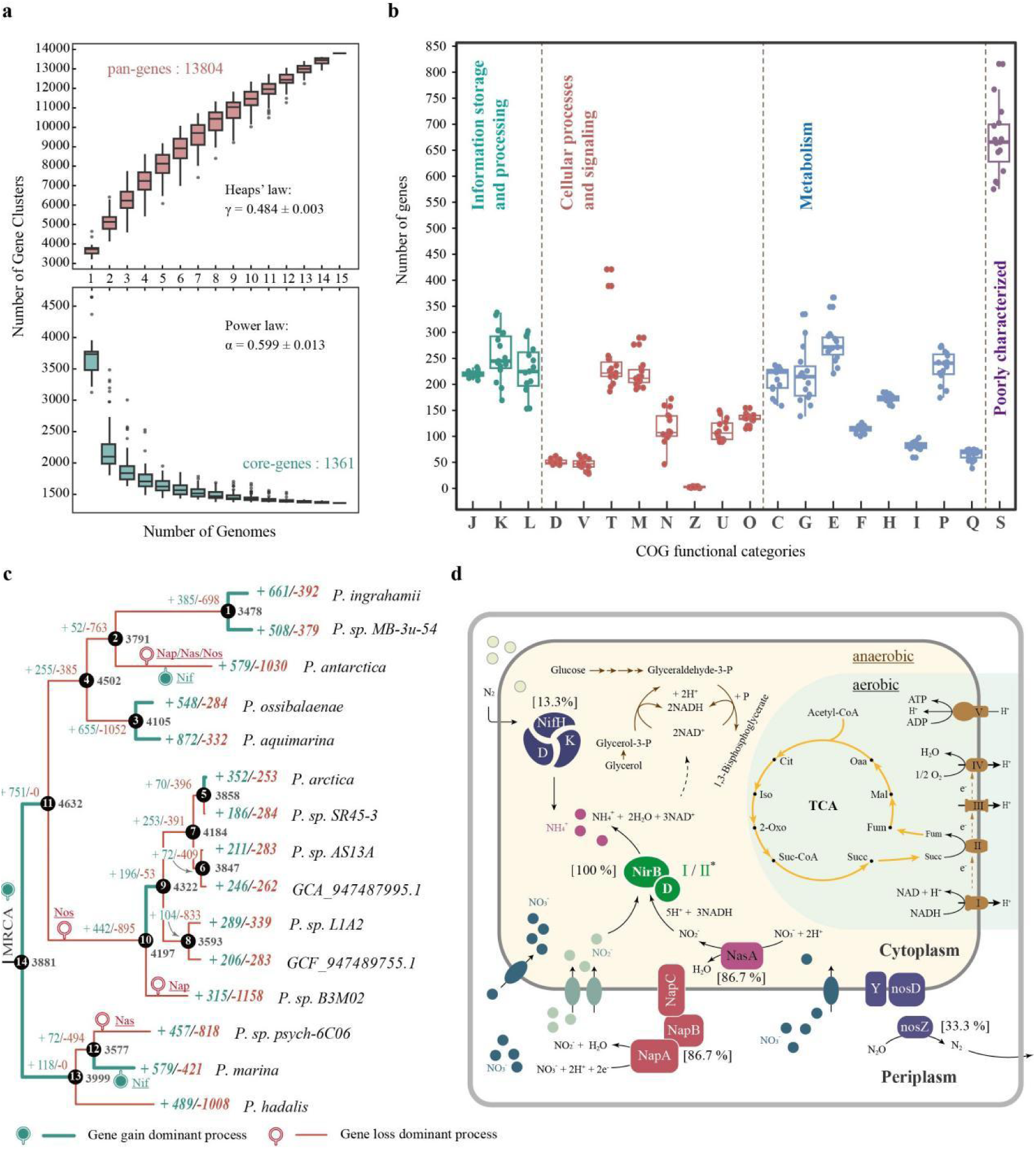
Insights into nitrogen-cycling genes from a pan-genomic analysis. **(a)** Rarefaction curves for pan and core genes in *Psychromonas*. Curves were fitted to the median values from 1,000 iterations using the Power Law model as previously described^50^. **(b)** COG functional analysis based on eggNOG-mapper. COG categories: J, translation, ribosomal structure and biogenesis; K, transcription; L, replication, recombination and repair; D, cell cycle control, cell division, chromosome partitioning; V, defense mechanisms; T, signal transduction mechanisms; M, cell wall/membrane/envelope biogenesis; N, cell motility; Z, cytoskeleton; U, intracellular trafficking, secretion and vesicular transport; O, post-translational modification, protein turnover and chaperones; C, energy production and conversion; G, carbohydrate transport and metabolism; E, amino acid transport and metabolism; F, nucleotide transport and metabolism; H, coenzyme transport and metabolism; I, lipid transport and metabolism; P, inorganic ion transport and metabolism; Q, secondary metabolites biosynthesis, transport and catabolism; S, function unknown. **(c)** Phylogenetic distribution of gene family gains and losses across *Psychromonas* lineages. The cladogram shows gains (blue) and losses (red) of gene families along each branch, for numbered internal nodes (1-14) and terminal strains. Solid blue circles indicate gene gain events associated with the N-cycle, open red circles indicate gene loss events associated with the N-cycle, and MRCA denotes the most recent common ancestor. **(d)** Overview of N-cycle relevant pathways in the *Psychromonas* genomes. Shapes indicate protein complexes or pathways present in the *Psychromonas* genomes. Numbers in square brackets indicate the percentage of genomes in which each protein is present. Elliptical symbols with arrows represent nitrate or nitrite transporters, and solid circles of different colours denote different forms of nitrogen. Roman numerals in the shapes representing electron transport chain complexes indicate: I, NADH dehydrogenase; II, succinate dehydrogenase; III, bc1 complex; IV, cytochrome cbb3 oxidase; and V, ATP synthase.

To assess fluctuations in gene number within individual categories, we introduced the coefficient of variation (CV) to quantify dispersion patterns (**Supplementary Fig. 3**). Overall, more than half of the enriched COG categories showed highly concentrated gene distributions (CV<15%), indicating that the *Psychromonas* genome is highly stable.

Next, we examined the evolutionary trajectory of genomic capacity, paying particular attention to the influx nodes of nitrogen cycle genes. Gene gain and loss analysis was employed to infer the history of genetic exchange between the genome and its environment. The common ancestor of *Psychromonas* was inferred, and the numbers of gained and lost gene families were calculated for each evolutionary node (nodes 1-14) and branch (**Figure 2c**). On the whole, genome contraction represents the major evolutionary trend in *Psychromonas*, although its most recent common ancestor (MRCA) experienced a significant early gene gain event. Such genome reduction may confer a survival advantage in cold, nutrient-poor environments. In addition, localized gene gain events tended to occur near the terminal stages of speciation, suggesting that *Psychromonas* inhabits environments characterized by high heterogeneity and diverse selective pressures.

What remains of greater interest is the evolutionary origin of nitrogen cycle related genes and the nodes where they were lost. At the MRCA, *Psychromonas* already possessed nitrate reductases (Nap/Nas), NADH-dependent nitrite reductase (NirBD) and nitrous oxide reductase (Nos). As evolution proceeds, a loss of *nos* occurred between evolutionary nodes 11 and 10, after which the sub-clade containing *P. arctica* formed a relatively unified nitrogen metabolism pattern, converting nitrate to nitrite and subsequently to ammonium. During the speciation of *P. antarctica* and *P. marina*, acquisition of *nif* genes conferred the capacity for nitrogen fixation. By contrast, nitrate reductases exhibited partial or even complete loss in some lineages. Notably, the capacity to reduce nitrite to ammonium was consistently retained throughout the evolutionary history of the entire lineage, suggesting that this function is indispensable for the survival of *Psychromonas*.

To understand the significance of nitrogen cycling, particularly nitrate assimilation (NO_3_^-^ to NH_4_^+^), in the energy metabolism of *Psychromonas*, we constructed a related metabolic model (**Figure 2d**). Under aerobic conditions, as in most aerobic microorganisms, *Psychromonas* relies primarily on the TCA cycle to metabolize organic substrates, thereby utilizing the respiratory chain for electron transfer and energy generation. Unlike denitrifiers, which switch to denitrification for respiration under anaerobic conditions, *Psychromonas* relies on substrate-level phosphorylation to meet its energy supply. Anaerobic conditions also activated nitrogen cycling processes in *Psychromonas*. Approximately 86.7% of genomes encode the periplasmic nitrate reductase (Nap) system (**Supplementary Fig. 2**), whose core components (NapC, NapB, and NapA) transfer electrons via the chain NapC-NapB-NapA to reduce nitrate. A similar abundance was observed for the cytoplasmic assimilatory nitrate reductase NasA, which not only reduces nitrate but also consumes NADH, helping maintain redox homeostasis. The resulting nitrite must be further converted to ammonium for incorporation into organic matter, a process carried out by the enzyme NirB through the consumption of the reducing power NADH. Furthermore, 13.3% of the genomes harbor the nitrogenase structural gene *nifH* along with the accessory genes *nifDK,* suggesting a putative nitrogen fixation capacity. Although we detected the periplasmic nitrous oxide reductase NosZ, an enzyme typically involved in denitrification, the other genes essential for a complete denitrification pathway were absent from *Psychromonas* genomes. Most importantly, NirB serves as a key enzyme in the nitrogen cycle of *Psychromonas*, catalyzing the oxidation of NADH to NAD^+^. The regenerated NAD^+^ then couples with the EMP pathway, accepting electrons released during substrate catabolism, thereby facilitating both the assimilation of inorganic nitrogen and the maintenance of cytoplasmic redox homeostasis. Moreover, the genomic organization of *nirB* itself is distinctive. It exists as two isoforms in the vast majority of *Psychromonas* genomes, a rare configuration to our knowledge.

### The coexistence of divergent NIR isoforms is a feature of *Psychromonas*

The coexistence of two NIR isoforms within a single genome, rather than a simple gene duplication event, may represent a unique genomic feature of the *Psychromonas* genus. Graphically, multiple sequence alignments (MSAs) of representative NirB clearly revealed sequence divergence between the two isoforms (**Figure 3a**). NirB homologs from the same genome were assigned to two different clusters (Clusters I and II). Nevertheless, the annotated functional regions remained highly conserved. Specifically, 17 out of 36 residues in the FAD-binding region and 23 out of 33 in the NADH-binding site were conserved. Moreover, the cysteine residues responsible for Fe-S cluster binding were completely invariant across both clusters. These cofactors are essential for electron transfer during the reduction of nitrite to ammonium.

**Figure 3.**
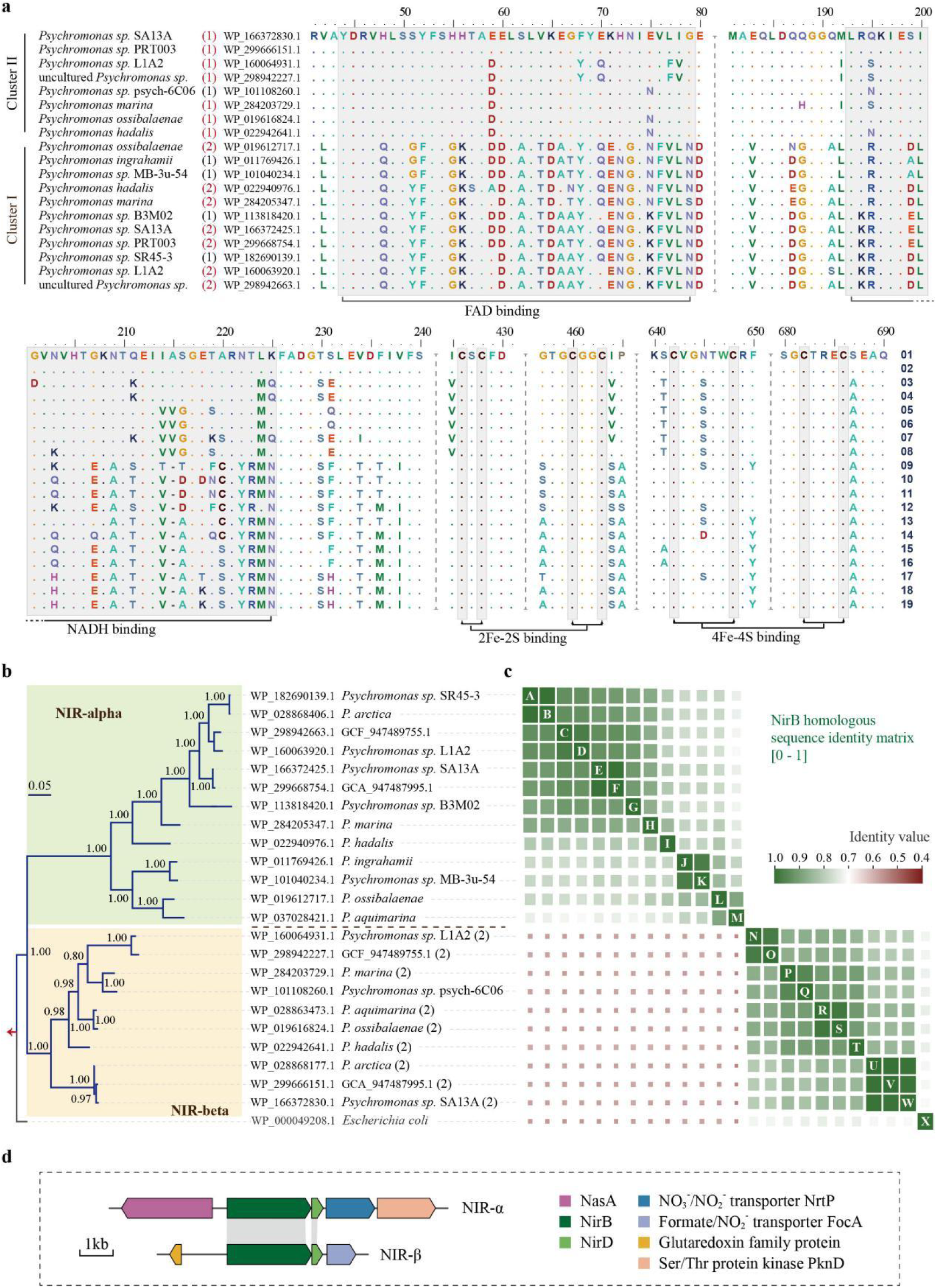
*Psychromonas* genomes contain two NIR isoforms. **(a)** Multiple sequence alignment of conserved regions in NirB proteins from *Psychromonas*. Sequence labels on the left comprise three components: species name, a parenthetical number and protein accession number. Red parenthetical numbers indicate genomes containing two NIR isoforms; the number denotes its occurrence count. In the alignment, a dot (.) denotes an amino acid identical to the reference residue in the first row of that column. **(b)** Phylogenetic tree of *Psychromonas* NirB protein sequences. *Escherichia coli* NirB was used as the outgroup by the method of Mrbayes. Tip labels show the protein accession number and species name. A parenthetical ‘(2)’ indicates genomes encoding two NIR isoforms. The clade highlighted in light green is designated the NIR-α group, and the clade highlighted in orange is the NIR-β group. Numbers at nodes represent Bayesian posterior probabilities (PP), with values ≥0.95 considered well supported. The red arrow indicates the root. **(c)** Heat map depicting pairwise sequence similarities, calculated as percent identity from all-against-all BLASTP alignments. Corresponding values are provided in Supplementary Table 6. Letter labels correspond to the sequence labels on the left. **(d)** The *nirB* operon and its flanking genes constitute a highly conserved genomic context within the genus *Psychromonas*.

To more accurately characterize the evolutionary divergence of NirB homologs, a Bayesian phylogenetic tree was constructed using the encoded protein sequences from all available genomes (**Figure 3b**). The tree clearly separates homologs into two clusters, here designated NIR-α and NIR-β for convenience in subsequent analyses. The NIR-α group comprises fourteen proteins and is present in all genomes except *Psychromonas* sp. psych-6C06, which contains only a single NIR-β system. All of the remaining genomes harboring NIR-β encode two NADH-dependent nitrite reductases, including *P. marina*, *P. aquimarina*, *P. ossibalaemae*, *P. hadalis*, *P. arctica*, *Psychromonas* sp. L1A2, *Psychromonas* sp. SA13A, and two uncultured genomes GCF_947489755.1 and GCA_947487995.1. These results suggest that a dual NIR system composed of NIR-α and NIR-β is the major genotype, although sole NIR-α (*Psychromonas* sp. SR45-3, *Psychromonas* sp. B3M02, *P. ingrahamii*, and *Psychromonas* sp. MB-3u-54) or sole NIR-β (*Psychromonas* sp. psych-6C06) is the minor genotype.

To better visualize the differences in sequence similarity among clusters, a pairwise sequence identity matrix was generated using the BLAST algorithm (**Figure 3c and Supplementary Table 6**). Sequence identity between the NIR-α and NIR-β groups was below 0.6, yet such a degree of divergence is rare among housekeeping genes within a genus. Intra-group sequence identity was generally above 0.8; in particular, some sequence pairs (e.g., AB and CD) showed even closer homology, reflecting the phylogenetic relatedness of their corresponding genomes. Together, the sequence identity matrix and phylogenetic tree provide mutually corroborating evidence, confirming the coexistence of two NIR lineages with distinct evolutionary rates in *Psychromonas*.

Additionally, the divergent genomic neighborhoods of *nirB* genes provide further support for the above conclusion. To identify conserved syntenic blocks, we conducted an analysis centered on the *nirB* gene, extracting and comparing sequences from 10 kb upstream and downstream. The flanking genes of *nirB* in different groups showed distinct syntenic patterns (**Figure 3d and Supplementary Table 7**). Conserved genes in the NIR-α group included the cytoplasmic assimilatory nitrate reductase *nasA*, the nitrate/nitrite transporter *nrtP* (also known as *narK*), and the serine/threonine-protein kinase *pknD*. As an important component of signal transduction, PknD may be involved in regulating expression of the NIR-α gene cluster. The conserved gene composition of the NIR-β group was relatively simple: *nirD* was adjacent to *focA*, which encodes a formate/nitrite transporter; and a conserved gene located further upstream of *nirB*, annotated as a glutaredoxin family protein. Although NirB-coding genes have different genetic surroundings in different NIR groups, they are typically located near nitrite transporter systems. Overall, this analysis shows that two isoforms of NADH-dependent nitrite reductase and their corresponding catalytic systems are widely present in *Psychromonas* genomes, and we find that they have distinct evolutionary rates, representing a characteristic genomic feature.

### Molecular basis for the differential evolution of two isoforms

The NADH-dependent nitrite reductase complex comprises large and small subunits, the former of which incorporates the essential cofactors FAD, NADH, [2Fe-2S] and [4Fe-4S] clusters, and siroheme. For a long time, the absence of structural information has limited a comprehensive understanding of the catalytic mechanism and functional principles of this complex molecular machine. To address this issue, we predicted the protein structure of nitrite reductase based on a large-scale distillation dataset (Method section).

From a global perspective, the large subunit NirB exhibits a clam-shaped architecture composed of two nearly symmetrical domains. The N- and C-terminal regions form two shell-like substructures with a flexible central region, imparting high conformational plasticity to the protein (**Figure 4a**). The large subunit relies on multiple cofactors to shuttle electrons from nitrite to ammonium. Given the critical role of the residues that coordinate these cofactors, we therefore mapped the specific interactions between each cofactor and its binding amino acids (**Figure 4b**). Our analysis confirmed the canonical cysteine coordination motif for the Fe-S cluster. The sulfur atoms of four cysteine residues (Cys641, Cys647, Cys681 and Cys685) serve as metal-sulfur ligands to the iron ions. This arrangement forms an efficient electron transfer pathway from the protein surface to the core of the [4Fe-4S] cluster. In contrast to the relatively simple, near-tetrahedral geometry of the [4Fe-4S] cluster-binding site, the FAD-binding pocket involves a network of 17 amino acid residues, constituting a far more intricate binding interface. Val14 and Val81 provide a stable hydrophobic core that accommodates the purine ring of FAD. Additionally, a hydrogen bond is formed with the side-chain hydroxyl group of Thr108. This interaction pattern is also observed in flavin-dependent thymidylate synthase (PDB ID: 3AH5)^54^. Furthermore, the negatively charged pyrophosphate chain of FAD is neutralized by the guanidinium groups of Arg45 and Arg129, forming strong salt bridges that further stabilize the cofactor conformation. Structurally, the C4 atom of the NADH nicotinamide ring and the N5 atom of the FAD isoalloxazine ring are approximately 3.5Å , and the two rings are nearly coplanar (**Supplementary Fig. 4**). This geometry is ideal for direct hydride ion transfer. In this configuration, the adenosine ring of NADH is anchored within the hydrophobic pocket formed by Ile150, Pro115, and Phe175, and its phosphate backbone is secured by the glycine-rich loop like motif comprising Gly151 and Gly153.

**Figure 4.**
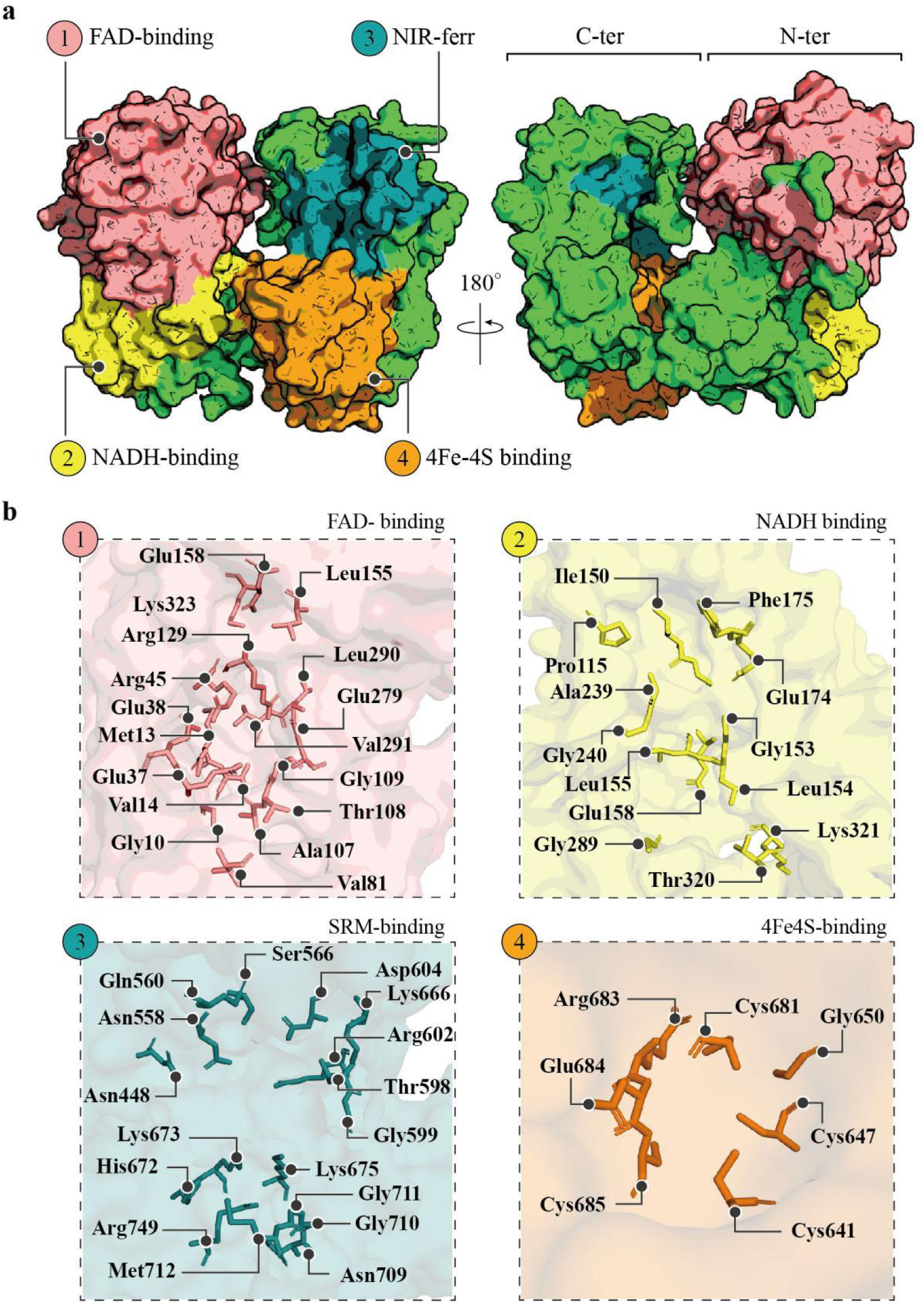
Structural and ligand-binding analysis of NADH-dependent nitrite reductase. **(a)** Structural representation of the NADH-dependent nitrite reductase larger subunit. The model is colour-coded by functional domain. **(b)** Zoomed-in view of the protein-ligand interaction interface of NirB. Amino-acid residues are shown in stick representation.

Subsequently, our analysis revealed that the siroheme-binding site is enriched in polar residues (13 out of 17), including Ser566, Asn448, and Thr598. Notably, the charged residues within this site are predominantly basic (e.g., Arg602, Lys666, His672), which likely facilitates interaction with the negatively charged substrate. An extensive hydrogen-bonding network formed by the positively charged residues and siroheme carboxylates serves to polarize the protonation state of the anionic substrate, thus enabling the efficient six-electron reduction of nitrite to ammonium. Electrostatic potential mapping of the surface further revealed the distinct electrostatic character of the catalytic pocket (**Figure 5a and b**). This region may act as a trap that draws NO ^-^into the pocket while limiting its escape and promotes release of the positively charged product NH_4_^+^. Structural details showed that, in addition to interacting with siroheme, nitrite forms direct contacts with Arg635, Lys672 and Lys674 at the center of this trap (**Figure 5c**). These three residues form a protrusion at the base of the positively charged trap, resembling a scaffold that support electron transfer between the substrate nitrite and siroheme. The same catalytic mechanism has also been confirmed in sulfite reductase, which catalyzes the analogous six-electron reduction of sulfite to sulfide^55^. In addition, we also investigated the functional contribution of the small subunit to catalysis. NirD is a small protein (shorter than 110 amino acid residues) that binds the C-terminus of NirB (**Figure 6a and b**), specifically contacting α-helices 16-19, 21, and 26-27 (**Figure 6d**). Analysis of the interface residues indicated that association between the large and small subunits is mediated primarily by polar interactions (**Supplementary Fig. 5**). By calculating the electrostatic potential (ESP) of the contact surface, we identified two pairs of oppositely charged interaction regions at the interface (**Figure 6c**). Electrostatic attraction between oppositely charged residues promotes protein approach and association, and stabilizes complex formation. This interaction helps to stabilize the spatial architecture of the catalytic center of NirB. In detail, the active site comprises two opposing helical bundles (left: α-helices 16-18; right: helices 19, 21, and 26-27) (**Figure 6d**). This configuration can be envisioned as a pair of pliers, with the helical bundles representing mobile jaws. NirD acts as a structural lock, fixing the relative orientation of these jaws to achieve the catalytically competent state (**Figure 6e**).

**Figure 5.**
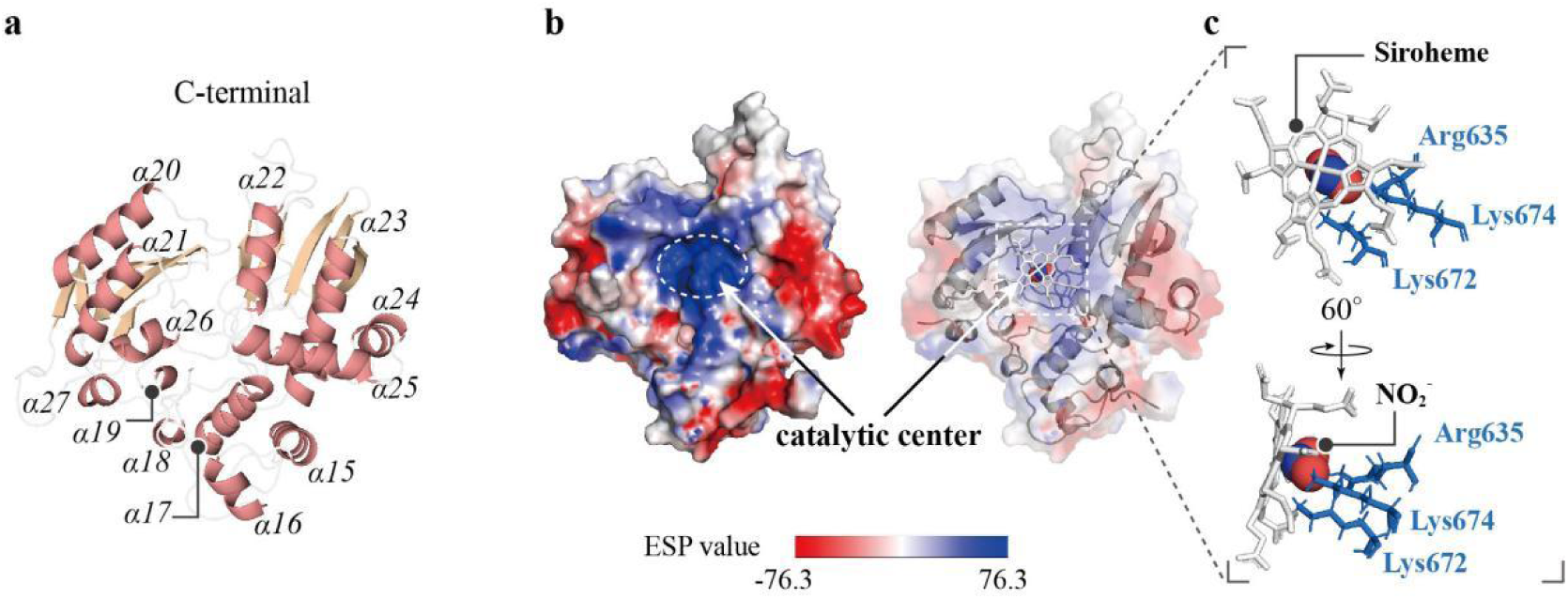
Structural basis for catalysis in NADH-dependent nitrite reductase. **(a)** Cartoon representation of the C-terminal structure of nitrite reductase. α-Helices are numbered sequentially according to their order in the amino acid sequence. **(b)** A positively charged catalytic core is revealed by electrostatic potential analysis combined with a transparent surface representation of the large-subunit active site. Siroheme is shown as white sticks. **(c)** Architecture of the catalytic center, with a close-up highlighting key amino acid residues involved in catalysis.

**Figure 6.**
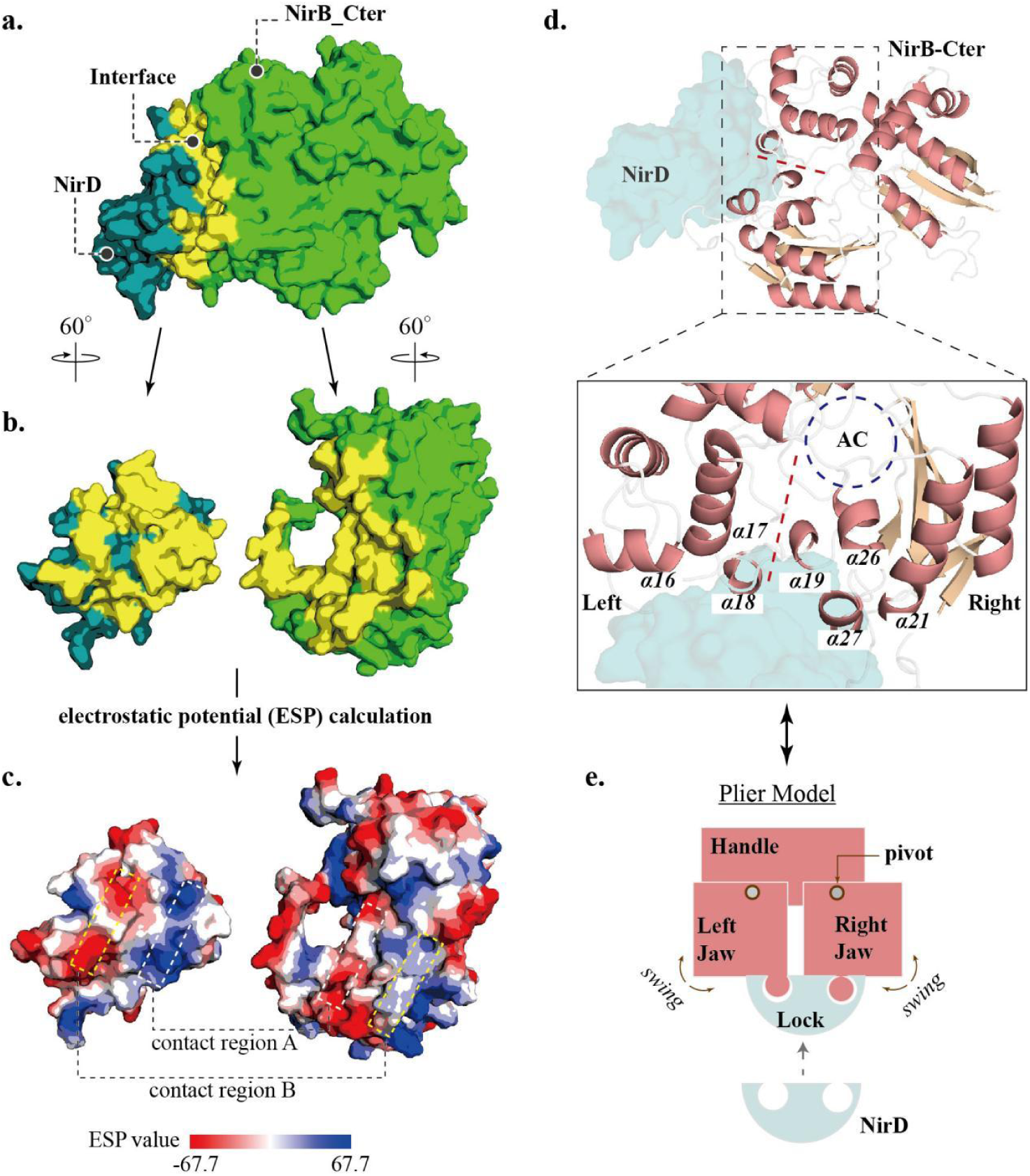
Assembly and functional mechanism of the nitrite reductase complex. **(a)** Structure of the NADH–dependent nitrite reductase heterodimer. The large subunit C–terminal domain is shown in lime green and the small subunit in blue; the interaction interface is highlighted in yellow. **(b)** Structural view highlighting the dimer interface. The two subunits are rotated by 60° about the y–axis to expose the buried contact surface. **(c)** Electrostatic complementarity at the interface. Surface electrostatic potential (ESP) of the separated subunits reveals two major contact regions (A and B) with complementary charges, indicating favorable electrostatic interactions. **(d)** NirD stabilizes the catalytic core of the large subunit. The arrangement resembles a pair of pliers: the left jaw (α–helices 16-18) and the right jaw (α–helices 19, 21, 26, and 27) are secured in a lock–like manner by NirD. A close-up view illustrates the large/small subunit interface that stabilizes the active-site conformation; numbers indicate α-helices, and the catalytic center is marked by a dashed black circle. **(e)** Plier-like model of the catalytic center.

Based on the aforementioned model, we compared the two NIR isoforms. Their C-terminal regions, which form the structural scaffold of the active site, are highly conserved (RMSD=0.67 Å). The small deviation is primarily due to the C-terminus of NIR-β folding into an α-helix over the last seven residues, whereas the corresponding segment in NIR-α remains unstructured (**Supplementary Fig. 6**). In contrast, the N-terminal regions of the two isoforms exhibit pronounced structural divergence. Although they share the same topological fold, rigid-body displacements are evident in several segments. For instance, α-helices 12-14 in both isoforms are nearly parallel in orientation, yet their axes are offset (**Supplementary Fig. 6 and 7**). These structural distinctions may underlie differences in their ligand-binding capabilities. Overall, our structural data elucidate the nitrite reductase mechanism and clarify the architectural similarities and differences between the two isoforms.

### Dual isoforms confer an adaptive advantage in *Psychromonas*

To investigate the potential biological advantage conferred by the coexistence of two NIR isoforms, we simulated culture conditions with different nitrogen sources to profile their expression patterns. We initially examined whether the expression of the two isoforms is induced by nitrite under anaerobic conditions. Mid-logarithmic cultures (OD_600_=0.5) (**Supplementary Fig. 8**) were incubated at 15℃ under anaerobic conditions in the presence or absence of 2 mM nitrite to assess the response of the nitrite reductase system to its substrate. Both isoforms showed strong induction by nitrite, with expression rising from nearly undetectable levels to a greater than 10-fold increase (**Figure 7a**). Furthermore, NIR-β expression modestly exceeded that of NIR-α following identical induction. However, it remains unclear whether this expression difference stems from positional effects within the operon or from distinct regulatory patterns governing the two isoforms. Accordingly, we established a time-course monitoring system to profile the dynamic expression of the *nirB* genes following induction (**Figure 7b**).

**Figure 7.**
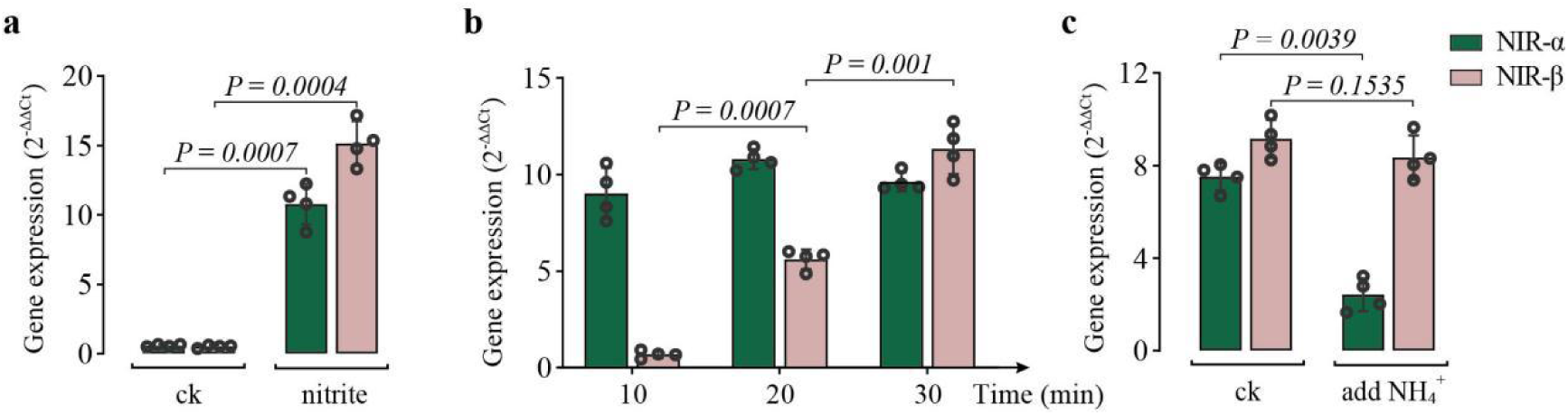
Comparative expression of the two *Nir* genes under different conditions. **(a)** Real-time quantitative PCR analysis of the two *Nir* isoforms following induction with 2 mM nitrite for 30 min. **(b)** Time-course analysis of *Nir* gene expression with nitrate as the sole nitrogen source. **(c)** Differential expression of the two *Nir* isoforms in response to NH_4_^+^. Dark green indicates NIR-α, and pink indicates NIR-β. Open circles represent individual data points. Statistical significance was determined using a two-tailed Student’s t-test.

During the early stage of induction, a marked difference in expression was observed between the two systems: the mean expression level of NIR-α was 9.03, whereas that of NIR-β was 0.69, approximately 13-fold lower. As induction progressed to 20 min, a higher expression level was maintained by the NIR-α *nirB* gene relative to the NIR-β, but the difference had narrowed substantially. Thus, divergent expression dynamics were observed between the two NIR systems. The copy linked to NIR-α showed stable expression over time, in contrast to the steadily increasing transcript levels of the NIR-β-linked copy. By the final time point, the NIR-β-associated *nirB* gene surpassed its NIR-α counterpart, reaching a mean expression level of 11.32. Therefore, divergent regulatory strategies likely evolved in the two NIR systems, refining their temporal expression patterns for specific adaptive needs.

We further investigated the feedback repression by ammonium, the end product of nitrite reductase, on the expression of *nirB* genes from different genomic loci. This inquiry is grounded in the fact that feedback inhibition is a core strategy for maintaining bacterial homeostasis and resource efficiency. The test was performed by supplementing the cultures with 5 mM NH_4_Cl and monitoring subsequent changes in *nirB* transcript levels (**Figure 7c**). Following ammonium addition, expression of the NIR-α *nirB* gene was reduced to 32% of its initial level, declining from 7.51 to 2.42. In contrast, NIR-β expression remained stable in the presence of ammonium. The two distinct expression modes in response to ammonium thus allow *Psychromonas* to maximize nitrite assimilation, an adaptation that may be critical for survival in cold and oligotrophic aquatic environments. Overall, the two-NIR genomic architecture not only facilitates a rapid response to environmental nitrate but also maximizes the utilization of available nitrogen resources, thereby enhancing adaptive fitness.

## Discussion

The significant role of *Psychromonas* in environmental nitrogen cycling has long been underappreciated, likely overshadowed by its dominant and more conspicuous functions in carbon cycle. Here, we gained new insights into the evolutionary history of *Psychromonas* and revealed specific genomic features associated with nitrogen cycling. This genus plays a key role in mediating the fixation of environmental dissolved inorganic nitrogen in cold habitats, thus driving microbial nitrogen immobilization. In particular, the products of both dissimilatory and assimilatory nitrate reduction converge on assimilatory nitrite reduction, and the pathway achieving nitrite assimilation is stably retained across *Psychromonas* genomes. Remarkably, a key genomic characteristic of *Psychromonas* is the coexistence of dual NIR systems, which broadens the scope of known nitrogen cycling strategies and confers a competitive advantage in cold nitrite assimilation.

*Psychromonas* can survive in low-temperature environments and even in the abyssal zone, which is characterized by a limited oxygen supply and often an uncertain availability of suitable terminal electron acceptors for respiration. Nonetheless, the glycolytic pathway sustains core metabolism by generating ATP and regenerating redox carriers NADH, thus maintaining metabolic flux. This results in an excess of reducing equivalents in the cytoplasm. Here, the NirBD complex consumes NADH and regenerates NAD^+^, thus maintaining cellular NADH/NAD^+^ homeostasis (**Figure 2d**). The NirBD system, which also requires anaerobic conditions for activation, catalyzes the reduction of nitrite to ammonium^56^. This reaction consumes five protons, thereby mitigating intracellular acidification that would otherwise result from proton accumulation. Additionally, the ammonium produced contributes to the maintenance of intracellular pH homeostasis^57^. Moreover, this reaction oxidizes three molecules of NADH per molecule of NH_4_^+^ produced, which is more efficient than NADH oxidation via NADH:ubiquinone oxidoreductase in respiratory complex I during aerobic respiration^58^. Although NirBD is not directly membrane-associated like NrfA, it likely contributes significantly to energy metabolism under oxygen limitation in *Psychromonas*^59^. Strong support comes from the work of Wang *et al.*, who demonstrated in *E. coli* that nitrite reduction to ammonium by NIR enhances cellular energy efficiency under anaerobic conditions, a mechanism likely conserved in *Psychromonas*^60^.

In response to persistent environmental challenges, bacteria have continuously evolved adaptive mechanisms to ensure survival and fitness. A key strategy is the evolution of isoenzyme systems, which confers adaptive advantage by providing both functional redundancy and specialization. The coexistence of two NAP isoforms in *Shewanella* exemplifies this strategy. This genus is an ideal model as it is well-studied and its members are often psychrophilic and barophilic^61^. Two NAP isoforms (NAP-α and NAP-β) confer a survival advantage in extreme environments through synergistic mechanisms^62^. First, their expression is complementarily regulated, with cold-induced NAP-α activity compensating for lower NAP-β levels under cold stress. Second, their distinct operon-encoded transporters synergistically enhance electron transport efficiency for nitrate ammonification. Together, these adaptations significantly enhance *Shewanella* fitness in extreme environments. Similarly, the thermophilic bacterium *Thermus* demonstrates this strategy. Our earlier work identified a case of isozyme coexistence within the denitrification pathway of *T. antranikianii*. Although they employ distinct metallo-cofactors, both the copper-containing and cytochrome cd₁ nitrite reductases catalyze the reduction of nitrite to nitric oxide. These two isoforms are differentially regulated upon oxygen limitation, facilitating a seamless transition in respiratory mode^63^. This metabolic plasticity is key to maintaining redox and energetic homeostasis within the high-temperature niche.

Here, we report that the dual NIR system in *Psychromonas* represents a novel genomic configuration of key bacterial nitrogen-cycling genes. Despite its organizational similarity to the *Shewanella* dual NAP system, the *Psychromonas* NIR homologs show greater divergence between two isoforms. Furthermore, the two systems may confer greater metabolic flexibility, allowing *Psychromonas* to optimize nitrate utilization and assimilation under fluctuating environmental parameters. Specifically, the two isoforms adopt specialized roles (**Figure 8**): NIR-α operates in a rapid-response mode to counter sudden nitrate surges, whereas NIR-β, being transcriptionally insensitive to ammonium, maintains constant production. This cooperative strategy maximizes nitrogen utilization in nutrient-poor environments.

**Figure 8.**
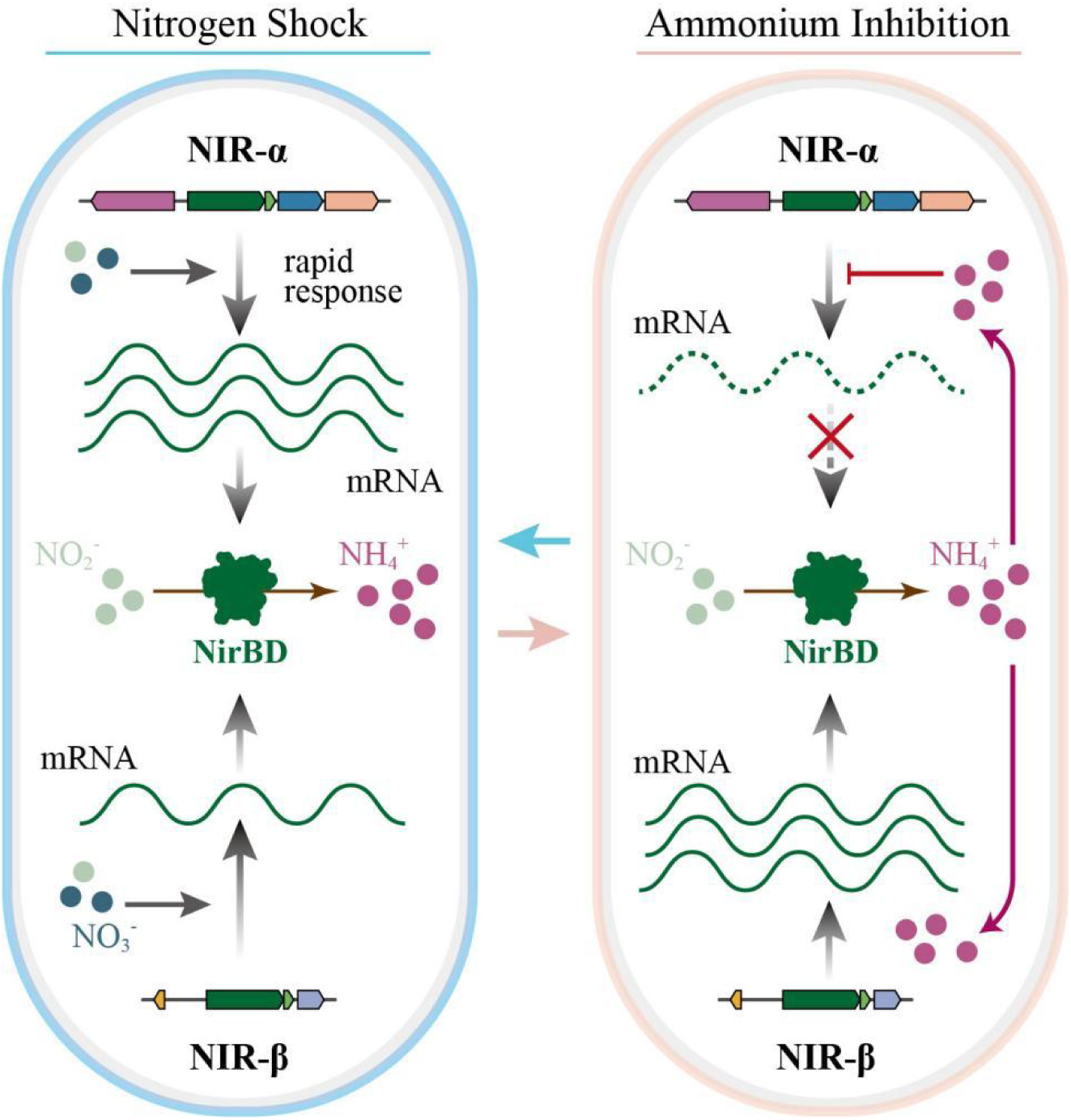
Molecular mechanisms of the dual NIR system underpin adaptation. Complementary transcription patterns of the NIR isoforms form a synergistic strategy. (Left) NIR-α responds rapidly to nitrate/nitrite, providing an advantage during nitrogen shock. (Right) NIR-β expression is insensitive to ammonium, allowing stable operation when NIR-α is feedback-inhibited, thereby maximizing dissolved inorganic nitrogen utilization.

Furthermore, the predicated protein structures provide critical structural insights, thereby facilitating an in-depth analysis of the molecular characteristics of the NIR isoforms. Our analysis reveals the intrinsic organization and stabilization mechanism of the NADH-dependent nitrite reductase active site, which explains how the enzyme retains functionality even in the absence of its small subunit (NirD)^64^. The mapping of multiple cofactor-binding sites has enriched the structural annotation of NirB. The conservation of these features, as corroborated by sulfite reductase structures, provided a basis for detailed comparison^65^. This analysis revealed structural divergence at the NADH-binding site between the two isoforms (**Supplementary Fig. 9**), differences that are likely to influence their respective catalytic activities.

While this study provides a systematic evaluation of the nitrogen-cycling potential in *Psychromonas* and a detailed characterization of the key NIR isoforms, it is limited by a genome dataset that does not cover all known species, which could lead to an underestimation of the genus’s genetic diversity.

In summary, this study re-evaluates the nitrogen cycling potential of *Psychromonas* in cold environments, demonstrating its significant role in microbial nitrogen immobilization and its broader ecological function. The unique dual NIR system not only enhances the adaptive fitness of *Psychromonas* but also provides a new perspective for understanding global biogeochemical cycles.We believe that continued exploration of microbial genomics will reveal additional novel strategies and mechanisms by which organisms adapt to their environments.

## Methods

### Dataset selection

A set of non-redundant *Psychromonas* genomes was downloaded from the National Center for Biotechnology Information (NCBI) RefSeq genome repository via the FTP site. Genome quality was assessed using CheckM v2^66^. High-quality genomes, defined as having completeness ≥95% and contamination <5% according to the recommendations of Bowers *et al.*^67^, were retained for subsequent analyses. Detailed genome information is provided in **Supplementary Table 1**.

### Construction of the phylogenetic tree

GTDB-Tk v2.3.2 was used to generate multiple sequence alignments (MSAs) of 120 bacterial marker genes^68^. A maximum-likelihood tree based on these MSAs was then constructed using IQ-TREE v2.4.0, with the best-fit model determined by ModelFinder to be LG+F+R4^69^.

For the phylogenetic analysis of NirB, all NirB proteins were extracted from *Psychromonas* genomes. Sequence alignment was performed using MUSCLE 5.1 with default parameters^70^. Gaps in the MSAs were trimmed using the automated mode of Trimal v1.4.rev15^71^. A Bayesian tree was then inferred using MrBayes 3.2.7a^72^ with the following parameters: ngen=1,000,000, nruns=2, nchains=4, diagnfreq=1,000, relburnin=yes, burninfrac=0.25, samplefreq=100, and printfreq=100. All trees were visualized and annotated using the R package ggtree v3.22^73^.

### Pan-genome analysis and functional annotation

Genomic data that passed the quality-control criteria were used for subsequent pan-genome analysis. Genome annotation was performed using Prokka v1.14.6^74^. Pan-genome ortholog clustering was conducted using PEPPAN v1.0.6^75^ with default parameters, including thresholds for core gene assignment (98%) and BLASTP identity (95%). The R package Micropan v2.1 was used to estimate whether the species had an open or closed pan-genome based on Heaps’ law (n=κN^γ^), using 1,000 permutations and 100 repeats^76, 77^. Gene cluster annotation was performed using eggNOG-mapper v2.1.6 and the eggNOG database v5.0^78, 79^.

### ANI and AAI calculations

To determine genomic species groups, we calculated pairwise ANI (average nucleotide identity) and AAI (average amino acid identity) values across the entire genome dataset (**Supplementary Table 3**). ANI values were calculated using fastANI v1.34.2, and AAI values were calculated using CompareM (https://github.com/donovan-h-parks/CompareM)^80^. A species boundary was defined as ANI >95% or AAI >95%; genomes exceeding these thresholds were considered to belong to the same species^81^.

### Gene gain and loss analysis

OrthoFinder v3.0.1b1 was used to generate a gene family count matrix for *Psychromonas*^82^. The *Psychromonas* genome tree was derived from the maximum-likelihood tree constructed by IQ-TREE. Gene gain and loss events during evolution were inferred using Count v25-beta, with Wagner parsimony and posterior probabilities as model settings^83^. The inferred results from Count were plotted using the R package ggtree^73^.

### Protein structural modelling

For structural predictions, SeedFold was adopted due to its excellent performance in both local (lDDT) and global (RMSD) accuracy assessments. For quality control, structures were also generated with AlphaFold for comparative assessment; the results further confirmed the reliability of the SeedFold models (**Supplementary Figure 10**)^84, 85^. The structural modelling of the NADH-dependent nitrite reductase large subunit was based on sequences WP_028868406.1 and WP_028868177.1, which were used to represent the NIR-α and NIR-β types, respectively. For the small subunit, the representative sequences were WP_028868407.1 and WP_028868176.1.

Protein structure alignments were performed using the online Pairwise Structure Alignment tool (https://www.rcsb.org/alignment), employing TM-align, jCE, jFATCAT and Smith-Waterman 3D as alignment methods. RMSD was used as the metric for evaluating structural alignment.

Protein-ligand complex modelling was performed using SeedFold-Linear, one of its features being the scaling of Pairformer width from 128 to 512 to increase model capacity^85^. For each complex, five structures were generated, and the model with the highest combined confidence score, pTM (predicted template modelling score) and ipTM (interface pTM) was selected for further analysis (**Supplementary Table 8**). All structural figures and protein-ligand interactions were visualized with PyMOL 3.1 (http://www.pymol.org).

### Bacterial strains and growth conditions

*P. marina* NBRC 103166 was used to investigate the expression of the two-NIR system under different conditions. Strains were cultured in marine 2216E medium at 15 ℃. A modified marine 2216E medium (Bacto peptone 5.0 g/L, yeast extract 1.0 g/L, NaCl 30.0 g/L, MgCl_2_ 5.9 g/L, Na_2_SO_4_ 3.24 g/L, CaCl_2_ 1.8 g/L, KCl 0.55 g/L) was used when growth under specific nitrogen sources was required and was supplemented with the corresponding nitrogen source as needed.

For anaerobic cultivation, medium without any electron acceptor was dispensed into 200-mL anaerobic culture bottles equipped with butyl rubber stoppers. A needle was inserted into the stopper during autoclaving and removed immediately after sterilization. Inorganic nitrogen salts were filter-sterilized in an anaerobic glove box and stored in anaerobic culture bottles. Bacterial growth curves were measured using BioTek Synergy 2 by monitoring absorbance at 600 nm.

### RNA extraction and reverse transcription

*P. marina* strains were cultured overnight at 15 ℃ and harvested at the exponential phase (OD_600_=0.5-0.6), then transferred to an anaerobic culture system containing a single nitrogen source. To stop transcription, 5% (v/v) phenol was added^86^. Cells were then collected by centrifugation at 12,000 rpm for 10 min at 4 ℃. Total RNA was extracted using the RNAprep Pure Cell/Bacteria Kit (Tiangen, Beijing, China) according to the manufacturer’s instructions. RNA quality was assessed using a NanoDrop 2000 spectrophotometer (Thermo Fisher Scientific). Extracted RNA samples were stored at -80 ℃ or used for subsequent experiments.

For cDNA synthesis, genomic DNA was removed from total RNA using gDNA wiper (Vazyme, Nanjing, China) at 42 ℃ for 2 min, followed by reverse transcription using the HiScript II 1st Strand cDNA Synthesis Kit (Vazyme, Nanjing, China) with random hexamers. cDNA products were stored at -20 ℃.

### Real-time qPCR

Real-time qPCR was modified according to the previously described method^87^, using SYBR Green I chemistry to quantify target cDNA on an Applied Biosystems 7500 instrument. The reaction mixture contained Taq Pro Universal SYBR qPCR Master Mix (Vazyme, Q712-02), and primers at a final concentration of 0.2 μM.

Relative expression levels of target genes were calculated using the 2^(-ΔΔCT) method described by Livak *et al.*^88^. *gyrB* was used as the internal control. Primers for real-time qPCR were designed based on target gene sequences from *P. marina* NBRC 103166, and primer pairs were selected using Primer-BLAST at NCBI (**Supplementary Table 9**).

## Statistical analysis and plots

All statistical analyses were performed using GraphPad Prism, and P values were calculated using a two-tailed Student’s t-test. All plot visualizations were generated using the R package ggplot2^89^.

## Data availability

Data supporting the findings of this study are available within the article and its Supplementary Information files. Original data are available in Mendeley Data (doi: 10.17632/4zvnd8stc4.1).

## Acknowledgments

This work was supported by the Guangdong Basic and Applied Basic Research Foundation [2021A1515110395 and 2022A1515011673].

## Author Contributions

Y.T. contributed to the conception and design of the study. Y.T. and R.L. drafted and revised the manuscript. Y.T. performed the physiological and biochemical experiments and the bioinformatic and statistical analyses. R.L. performed protein structure modeling, protein evolutionary analysis, and data analysis. All authors contributed to the manuscript and approved the submitted version.

## References

1. Feller, G. & Gerday, C. Psychrophilic enzymes: hot topics in cold adaptation. Nat. Rev. Microbiol. 1, 200–208 (2003).

2. Rodrigues, D. F. & Tiedje, J. M. Coping with our cold planet. Appl. Environ. Microbiol. 74, 1677–1686 (2008).

3. Staley, J. T. & Gosink, J. J. Poles apart: biodiversity and biogeography of sea ice bacteria. Annu. Rev. Microbiol. 53, 189–215 (1999).

4. Cavicchioli, R. Cold-adapted archaea. Nat. Rev. Microbiol. 4, 331–343 (2006).

5. Casanueva, A., Tuffin, M., Cary, C. & Cowan, D. A. Molecular adaptations to psychrophily: the impact of ‘omic’ technologies. Trends Microbiol. 18, 374–381 (2010).

6. Margesin, R. & Miteva, V. Diversity and ecology of psychrophilic microorganisms. Res. Microbiol. 162, 346–361 (2011).

7. Ramasamy, K. P. et al. Comprehensive insights on environmental adaptation strategies in Antarctic bacteria and biotechnological applications of cold adapted molecules. Front. Microbiol. 14, 1197797 (2023).

8. Cowan, D. A. Cryptic microbial communities in Antarctic deserts. Proc. Natl. Acad. Sci. USA 106, 19749–19750 (2009).

9. Allen, M. A. et al. The genome sequence of the psychrophilic archaeon, *Methanococcoides burtonii*: the role of genome evolution in cold adaptation. ISME J. 3, 1012–1035 (2009).

10. De Maayer, P., Anderson, D., Cary, C. & Cowan, D. A. Some like it cold: understanding the survival strategies of psychrophiles. EMBO Rep. 15, 508–517 (2014).

11. Lehnert, N., Dong, H. T., Harland, J. B., Hunt, A. P. & White, C .J. Reversing nitrogen fixation. Nat. Rev. Chem. 2, 278–289 (2018).

12. Sogin, M. L., et al. Microbial diversity in the deep sea and the underexplored “rare biosphere”. Proc. Natl. Acad. Sci. USA 103, 12115–12120 (2006).

13. Poff, K. E., Leu, A. O., Eppley, J. M., Karl, D. M. & DeLong, E. F. Microbial dynamics of elevated carbon flux in the open ocean’s abyss. Proc. Natl. Acad. Sci. USA 118, e2018269118 (2021).

14. Hutchins, D. A. & Capone, D. G. The marine nitrogen cycle: new developments and global change. Nat. Rev. Microbiol. 20, 401–414 (2022).

15. Zhang, X. et al. Microbial diversity and biogeochemical cycling potential in deep-sea sediments associated with seamount, trench, and cold seep ecosystems. Front. Microbiol. 13, 1029564 (2022).

16. Quan, Q., Liu, J., Li, C., Ke, Z. & Tan, Y. Insights into prokaryotic communities and their potential functions in biogeochemical cycles in cold seep. mSphere 9, e0054924 (2024).

17. Moran, M. A. & Durham, B. P. Sulfur metabolites in the pelagic ocean. Nat. Rev. Microbiol. 17, 665–678 (2019).

18. Bienhold, C., Boetius, A. & Ramette, A. The energy-diversity relationship of complex bacterial communities in Arctic deep-sea sediments. ISME J. 6, 724–732 (2012).

19. García-Lopez, E. et al. Microbial community structure driven by a volcanic gradient in glaciers of the Antarctic Archipelago South Shetland. Microorganisms 9, 392 (2021).

20. Doytchinov, V. V., Peykov, S. & Dimov, S. G. Study of the Bacterial, Fungal, and Archaeal Communities Structures near the Bulgarian Antarctic Research Base “St. Kliment Ohridski” on Livingston Island, Antarctica. Life (Basel) 14, 278 (2024).

21. Fernández-Gómez, B. et al. Bacterial community structure in a sympagic habitat expanding with global warming: brackish ice brine at 85-90 °N. ISME J. 13, 316–333 (2019).

22. Moncada, C. et al. Niche separation in bacterial communities and activities in porewater, loosely attached, and firmly attached fractions in permeable surface sediments. ISME J. 18, wrae159 (2024).

23. Fan, S. et al. Scientific and technological progress in the microbial exploration of the hadal zone. Mar. Life Sci. Technol. 4, 127–137 (2021).

24. Zhang, H. et al. The amphipod genome reveals population dynamics and adaptations to hadal environment. Cell 188, 1378–1392.e18 (2025).

25. Tamby, A., Sinninghe Damsté, J. S. & Villanueva, L. Microbial membrane lipid adaptations to high hydrostatic pressure in the marine environment. Front. Mol. Biosci. 9, 1058381 (2023).

26. Shah, A. M., Yang, W., Mohamed, H., Zhang, Y. & Song, Y. Microbes: a hidden treasure of polyunsaturated fatty acids. Front. Nutr. 9, 827837 (2022).

27. Methé, B. A., et al. The psychrophilic lifestyle as revealed by the genome sequence of *Colwellia psychrerythraea* 34H through genomic and proteomic analyses. Proc. Natl. Acad. Sci. USA 102, 10913–10918 (2005).

28. Metz, J. G. et al. Production of polyunsaturated fatty acids by polyketide synthases in both prokaryotes and eukaryotes. Science 293, 290–293 (2001).

29. León-Zayas, R. et al. Single cells within the Puerto Rico trench suggest hadal adaptation of microbial lineages. Appl. Environ. Microbiol. 81, 8265–8276 (2015).

30. Thomas, F. et al. Isotopic tracing reveals single-cell assimilation of a macroalgal polysaccharide by a few marine Flavobacteria and Gammaproteobacteria. ISME J. 15, 3062–3075 (2021).

31. Suominen, S., Dombrowski, N., Sinninghe Damsté, J. S. & Villanueva, L. A diverse uncultivated microbial community is responsible for organic matter degradation in the Black Sea sulphidic zone. Environ. Microbiol. 23, 2709–2728 (2021).

32. Zhang, W. et al. Genome reduction in *Psychromonas* species within the gut of an amphipod from the ocean’s deepest point. mSystems 3, e00009–18 (2018).

33. Hoffmann, K., Hassenrück, C., Salman-Carvalho, V., Holtappels, M. & Bienhold, C. Response of bacterial communities to different detritus compositions in Arctic deep-sea sediments. Front. Microbiol. 8, 266 (2017).

34. McDonald, N. D., Lubin, J. B., Chowdhury, N. & Boyd, E. F. Host-derived sialic acids are an important nutrient source required for optimal bacterial fitness in vivo. mBio 7, e02237–15 (2016).

35. Yu, X. A. et al. Low-level resource partitioning supports coexistence among functionally redundant bacteria during successional dynamics. ISME J. 18, wrad013 (2024).

36. Zhao, Y. et al. The origin, properties, structure, catalytic mechanism, and applications of fucoidan-degrading enzymes. Mar. Drugs 23, 97 (2025).

37. Vickers, C. et al. Endo-fucoidan hydrolases from glycoside hydrolase family 107 (GH107) display structural and mechanistic similarities to α-l-fucosidases from GH29. J. Biol. Chem. 293, 18296–18308 (2018).

38. Lozada, M., Diéguez, M. C., García, P. E. & Dionisi, H. M. Microbial communities associated with kelp detritus in temperate and subantarctic intertidal sediments. Sci. Total Environ. 857, 159392 (2023).

39. Kim, J., Kang, S., Kim, H. S., Kim, S. & Lee, S. S. Pilot plant study on nitrogen and phosphorus removal in marine wastewater by marine sediment with sequencing batch reactor. PLOS ONE 15, e0233042 (2020).

40. Jones, C. M., Graf, D. R., Bru, D., Philippot, L. & Hallin, S. The unaccounted yet abundant nitrous oxide-reducing microbial community: a potential nitrous oxide sink. ISME J. 7, 417–426 (2013).

41. Kim, D. D. et al. Identification of *nosZ*-expressing microorganisms consuming trace N2O in microaerobic chemostat consortia dominated by an uncultured *Burkholderiales*. ISME J. 16, 2087–2098 (2022).

42. León-Zayas, R. et al. Single cells within the Puerto Rico trench suggest hadal adaptation of microbial lineages. Appl. Environ. Microbiol. 81, 8265–8276 (2015).

43. Peoples, L. M. et al. A deep-sea isopod that consumes *Sargassum* sinking from the ocean’s surface. Proc. Biol. Sci. 291, 20240823 (2024).

44. Sun, H. et al. Novel insights into the rhizosphere and seawater microbiome of *Zostera marina* in diverse mariculture zones. Microbiome 12, 27 (2024).

45. Zehr, J. P. & Capone, D. G. Unsolved mysteries in marine nitrogen fixation. Trends Microbiol. 32, 532–545 (2024).

46. Nogi, Y., Hosoya, S., Kato, C. & Horikoshi, K. *Psychromonas hadalis* sp. nov., a novel piezophilic bacterium isolated from the bottom of the Japan Trench. Int. J. Syst. Evol. Microbiol. 57, 1360–1364 (2007).

47. Chun, J. et al. Proposed minimal standards for the use of genome data for the taxonomy of prokaryotes. Int. J. Syst. Evol. Microbiol. 68, 461–466 (2018).

48. Konstantinidis, K. T., Rosselló-Móra, R. & Amann, R. Uncultivated microbes in need of their own taxonomy. ISME J. 11, 2399–2406 (2017).

49. Jain, C., Rodriguez-R, L. M., Phillippy, A. M., Konstantinidis, K. T. & Aluru, S. High throughput ANI analysis of 90K prokaryotic genomes reveals clear species boundaries. Nat. Commun. 9, 1–8 (2018).

50. Tettelin, H., Riley, D., Cattuto, C. & Medini, D. Comparative genomics: the bacterial pan-genome. Curr. Opin. Microbiol. 11, 472–477 (2008).

51. Matthews, C. A., Watson-Haigh, N. S., Burton, R. A. & Sheppard, A. E. A gentle introduction to pangenomics. Brief Bioinform. 25, bbae588 (2024).

52. Abby, S. & Daubin, V. Comparative genomics and the evolution of prokaryotes. Trends Microbiol. 15, 135–141 (2007).

53. Pountain, A. W. et al. Transcription-replication interactions reveal bacterial genome regulation. Nature 626, 661–669 (2024).

54. Zhang, X. et al. Crystal structure of a flavin-dependent thymidylate synthase from *Helicobacter pylori* strain 26695. Protein Pept. Lett. 19, 1225–1230 (2012).

55. Smith, K. W. & Stroupe, M. E. Mutational analysis of sulfite reductase hemoprotein reveals the mechanism for coordinated electron and proton transfer. Biochemistry 51, 9857–9868 (2012).

56. Wang, H. & Gunsalus, R. P. The *nrfA* and *nirB* nitrite reductase operons in *Escherichia coli* are expressed differently in response to nitrate than to nitrite. J. Bacteriol. 182, 5813–5822 (2000).

57. Krulwich, T. A., Sachs, G. & Padan, E. Molecular aspects of bacterial pH sensing and homeostasis. Nat. Rev. Microbiol. 9, 330–343 (2011).

58. Berrisford, J. M., Baradaran, R. & Sazanov, L. A. Structure of bacterial respiratory complex I. Biochim. Biophys. Acta. 1857, 892–901 (2016).

59. Einsle, O., Messerschmidt, A., Huber, R., Kroneck, P. M. & Neese, F. Mechanism of the six-electron reduction of nitrite to ammonia by cytochrome c nitrite reductase. J. Am. Chem. Soc. 124, 11737–11745 (2002).

60. Wang, X. et al. The role of the NADH-dependent nitrite reductase, Nir, from *Escherichia coli* in fermentative ammonification. Arch. Microbiol. 201, 519–530 (2019).

61. Simpson, P. J. L., Richardson, D. J. & Codd, R. The periplasmic nitrate reductase in *Shewanella*: the resolution, distribution and functional implications of two NAP isoforms, NapEDABC and NapDAGHB. Microbiology (Reading) 156, 302–312 (2010).

62. Chen, Y., Wang, F., Xu, J., Mehmood, M. A. & Xiao, X. Physiological and evolutionary studies of NAP systems in *Shewanella piezotolerans* WP3. ISME J. 5, 843–855 (2011).

63. Liu, R. R. et al. Distinct expression of the two NO-forming nitrite reductases in *Thermus antranikianii* DSM 12462^T^ improved environmental adaptability. Microb. Ecol. 80, 614–626 (2020).

64. Yılmaz, H. et al. Nitrite is reduced by nitrite reductase NirB without small subunit NirD in *Escherichia coli*. J. Biosci. Bioeng. 134, 393–398 (2022).

65. Ghazi Esfahani, B., et al. Structure of dimerized assimilatory NADPH-dependent sulfite reductase reveals the minimal interface for diflavin reductase binding. Nat. Commun. 16, 2955 (2025).

66. Parks, D. H., Imelfort, M., Skennerton, C. T., Hugenholtz, P. & Tyson, G. W. CheckM: assessing the quality of microbial genomes recovered from isolates, single cells, and metagenomes. Genome Res. 25, 1043–1055 (2015).

67. Bowers, R. M. et al. Minimum information about a single amplified genome (MISAG) and a metagenome-assembled genome (MIMAG) of bacteria and archaea. Nat Biotechnol. 35, 725–731 (2017).

68. Chaumeil, P. A., Mussig, A. J., Hugenholtz, P. & Parks, D. H. GTDB-Tk v2: memory friendly classification with the genome taxonomy database. Bioinformatics 38, 5315–5316 (2022).

69. Minh, B. Q. et al. IQ-TREE 2: new models and efficient methods for phylogenetic inference in the genomic era. Mol. Biol. Evol. 37, 1530–1534 (2020).

70. Edgar, R. C. MUSCLE5: high-accuracy alignment ensembles enable unbiased assessments of sequence homology and phylogeny. Nat. Commun. 13, 6968 (2022).

71. Capella-Gutiérrez, S., Silla-Martínez, J. M. & Gabaldón, T. trimAl: a tool for automated alignment trimming in large-scale phylogenetic analyses. Bioinformatics 25, 1972–1973 (2009).

72. Ronquist, F. & Huelsenbeck, J. P. MRBAYES 3: Bayesian phylogenetic inference under mixed models. Bioinformatics 19, 1572–1574 (2003).

73. Xu, S., et al. ggtree: a serialized data object for visualization of a phylogenetic tree and annotation data. iMeta 1, e56 (2022).

74. Seemann, T. Prokka: rapid prokaryotic genome annotation. Bioinformatics 30, 2068–2069 (2014).

75. Zhou, Z., Charlesworth, J. & Achtman, M. Accurate reconstruction of bacterial pan- and core genomes with PEPPAN. Genome Res. 30, 1667–1679 (2020).

76. Snipen, L. & Liland, K. H. micropan: an R-package for microbial pan-genomics. BMC Bioinformatics 16, 79 (2015).

77. Tettelin, H., Riley, D., Cattuto, C. & Medini, D. Comparative genomics: the bacterial pan-genome. Curr. Opin. Microbiol. 11, 472–477 (2008).

78. Cantalapiedra, C. P., Hernández-Plaza A., Letunic, I., Bork, P., & Huerta-Cepas, J. eggNOG-mapper v2: functional annotation, orthology assignments, and domain prediction at the metagenomic scale. Mol. Biol. Evol. 38, 5825–5829 (2021).

79. Huerta-Cepas, J. et al. eggNOG 5.0: a hierarchical, functionally and phylogenetically annotated orthology resource based on 5090 organisms and 2502 viruses. Nucleic Acids Res. 47, D309–D314 (2019).

80. Jain, C., Rodriguez-R, L. M., Phillippy, A. M., Konstantinidis, K. T. & Aluru, S. High throughput ANI analysis of 90K prokaryotic genomes reveals clear species boundaries. Nat. Commun. 9, 5114 (2018).

81. Richter, M. & Rosselló-Móra, R. Shifting the genomic gold standard for the prokaryotic species definition. Proc. Natl Acad. Sci. USA 106, 19126–19131 (2009).

82. Emms, D. M. & Kelly, S. OrthoFinder: phylogenetic orthology inference for comparative genomics. Genome Biol. 20, 238 (2019).

83. Csűrös, M. Count: evolutionary analysis of phylogenetic profiles with parsimony and likelihood. Bioinformatics 26, 1910–1912 (2010).

84. Jumper, J. et al. Highly accurate protein structure prediction with AlphaFold. Nature 596, 583–589 (2021).

85. Zhou, Y., et al. SeedFold: scaling biomolecular structure prediction. *arXiv preprint* arXiv:2512.24354 (2025).

86. Owen, S. V. et al. A window into lysogeny: revealing temperate phage biology with transcriptomics. Microb. Genom. 6, e000330 (2020).

87. Liu, R. et al. An uncharacterized small protein MicN mediates the transcriptional reprogramming of *Salmonella* through regulating the RpoS-RNA polymerase interaction. *Commun*. Biol. 8, 1495 (2025).

88. Livak, K. J. & Schmittgen, T. D. Analysis of relative gene expression data using real-time quantitative PCR and the 2^(-ΔΔCT) method. Methods 25, 402–408 (2001).

89. Wickham, H. ggplot2: Elegant Graphics for Data Analysis. (Springer, 2016).

