## Supplementary Information for "Nitrite reductase NirB mediates an unconventional nitrogen assimilation strategy to enhance adaptations of extremophile"

Ruirui Liu and Ye Tian*

State Key Laboratory of Biocontrol, School of Life Sciences, Sun Yat-Sen University, Guangzhou, China

*Corresponding author: Ye Tian

### Supplementary Figure 1-10

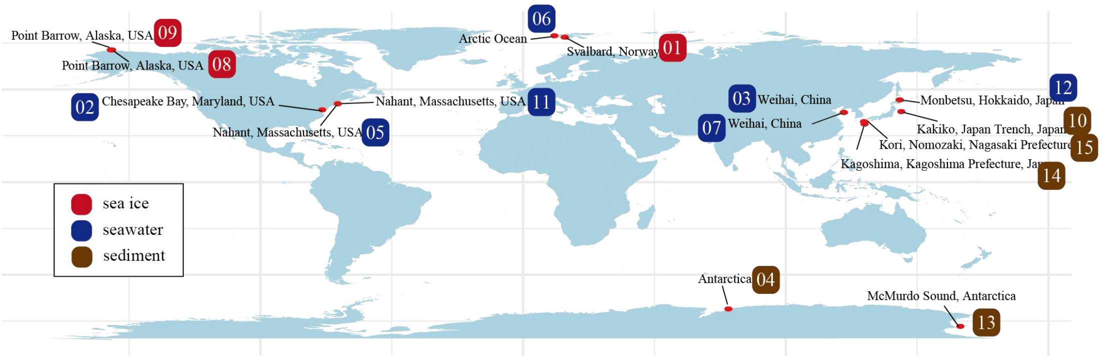


**Supplementary Fig. 1 Geographic distribution of Psychromonas genomic data.** To facilitate the interpretation of the sampling sites, their geographical locations are annotated. Detailed records for each site correspond to those listed in Supplementary Table 2. Boxes indicate the sampling locations of the selected genomes. Colors represent different sample types.

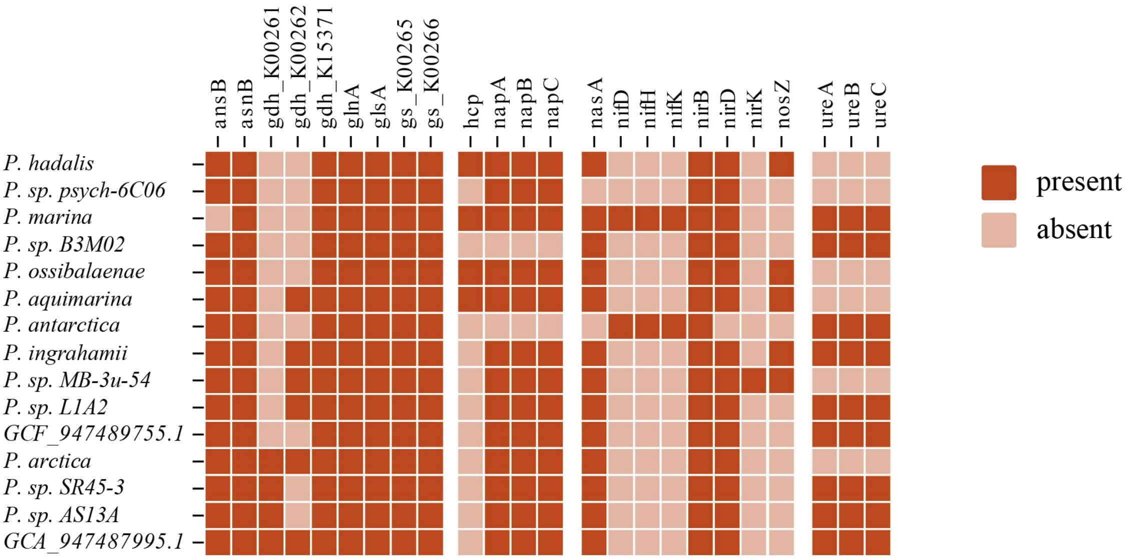


**Supplementary Fig. 2 Heatmap of N-cycle-related genes in *Psychromonas* genomes.** Presence (dark orange) and absence (light orange) of genes are indicated. Gene annotations beyond those described in the main text are provided below: *ansB*, Glutaminase-asparaginase; *asnB*, Asparagine synthase (glutamine-hydrolysing); *gdh_K00261*, Glutamate dehydrogenase (NAD(P)+); *gdh_K00262*, Glutamate dehydrogenase (NADP+); *gdh_K15371*, Glutamate dehydrogenase; *glnA*, Glutamine synthetase; *glsA*, Glutaminase; *gs_K00265*, Glutamate synthase (NADPH/NADH) large chain; *gs_K00266*, Glutamate synthase (NADPH/NADH) small chain; *ure*, Urease.

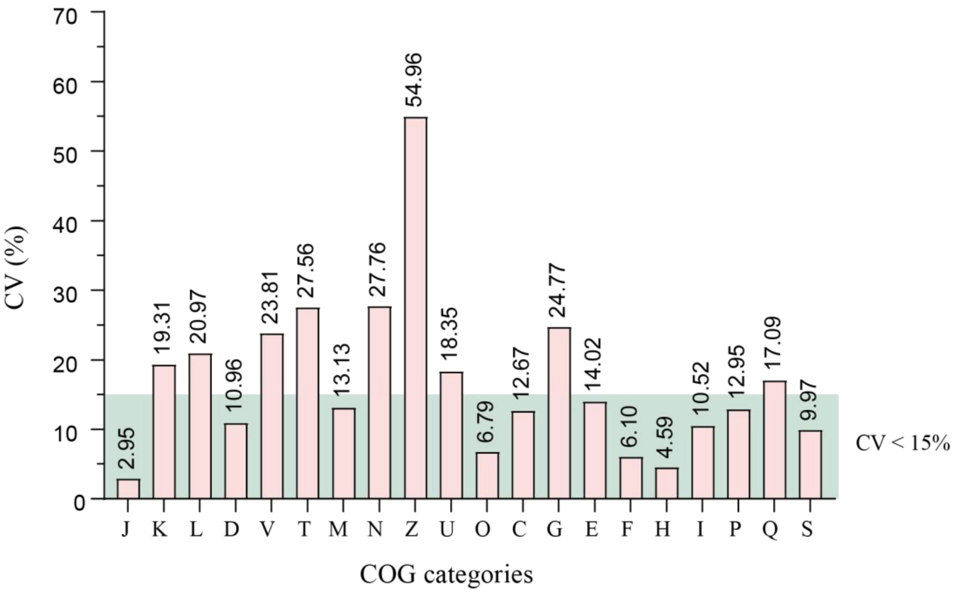


**Supplementary Fig. 3 Statistics of the coefficient of variation (CV) based on COG (Clusters of Orthologous Groups) annotations.** To assess fluctuations in gene number within individual categories, we introduced the coefficient of variation (CV) to quantify dispersion patterns. The letters on the x-axis denote COG categories; the numerical CV value is shown above each bar; The green shaded area indicates a CV of less than 15%. The calculation method is as follows: CV=(Standard Deviation/Mean)×100%. More than half of the enriched COG categories showed highly concentrated gene distributions (CV<15%), indicating that the *Psychromonas* genome is highly stable.

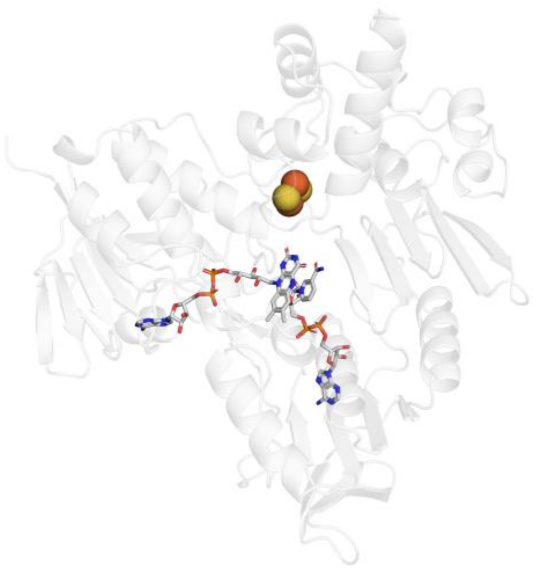

**Supplementary Fig. 4 The relative arrangement of NADH and FAD within the NirB protein.** The C4 atom of the NADH nicotinamide ring and the N5 atom of the FAD isoalloxazine ring are approximately 3.5Å , and the two rings are nearly coplanar.

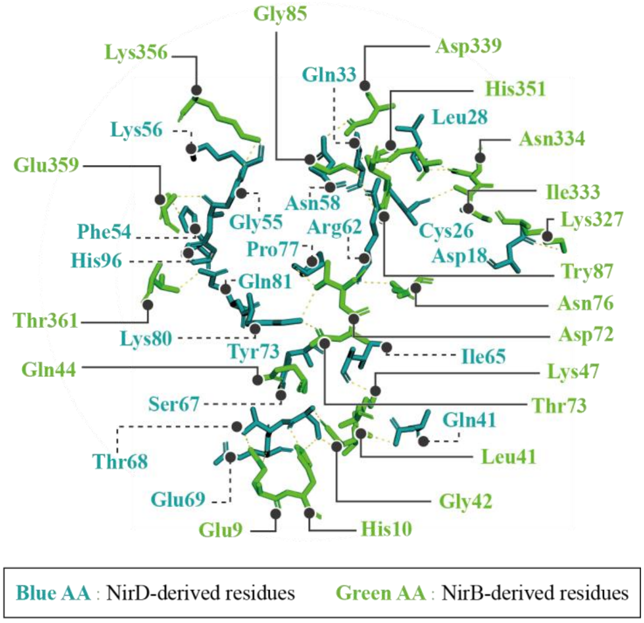


**Supplementary Fig. 5. Analysis of the interaction interface between the large (NirB) and small (NirD) subunits of NADH-dependent nitrite reductase.** Green residues are derived from NirB; blue residues are derived from NirD.

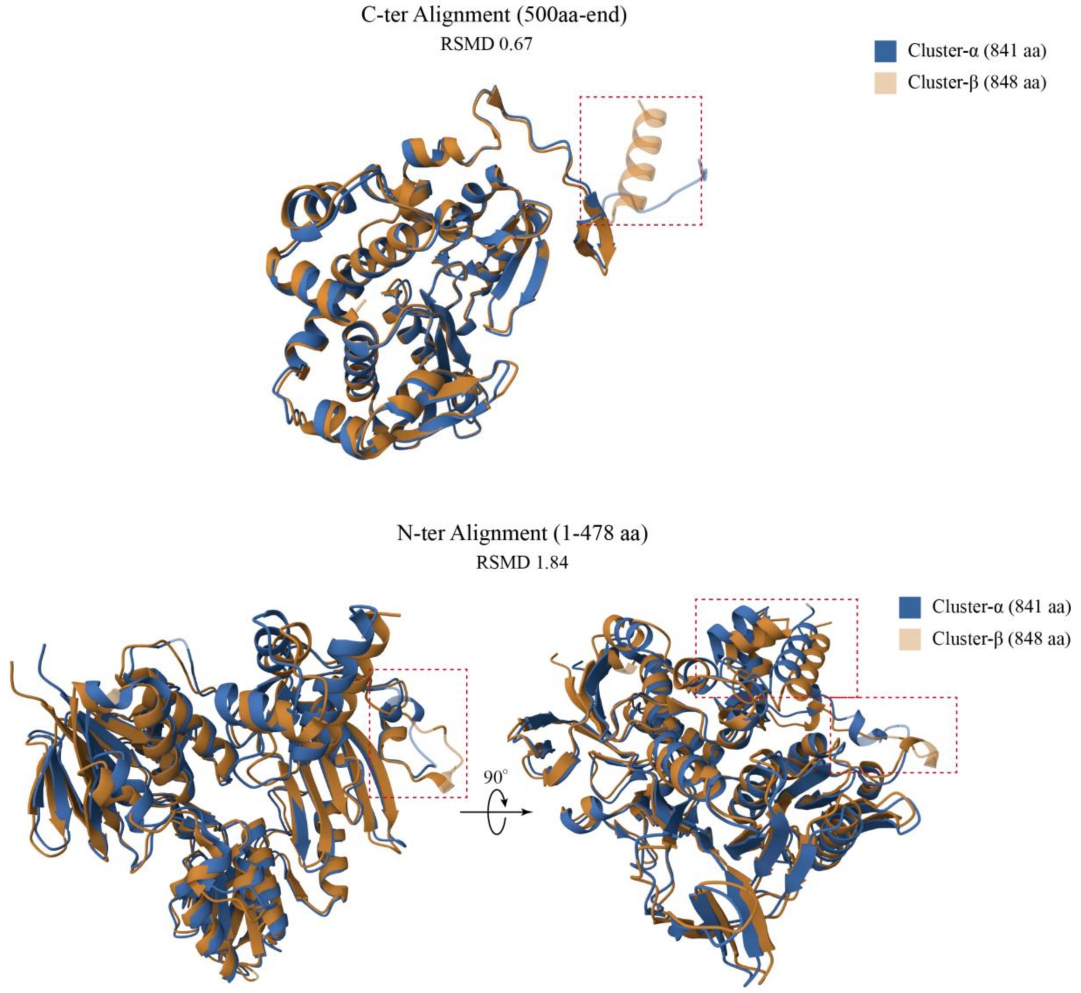


**Supplementary Fig. 6 Structural alignment of the two NIR isoforms.** The α-isoform is shown in blue cartoon, the β-isoform in orange. Regions of structural divergence are boxed in red.

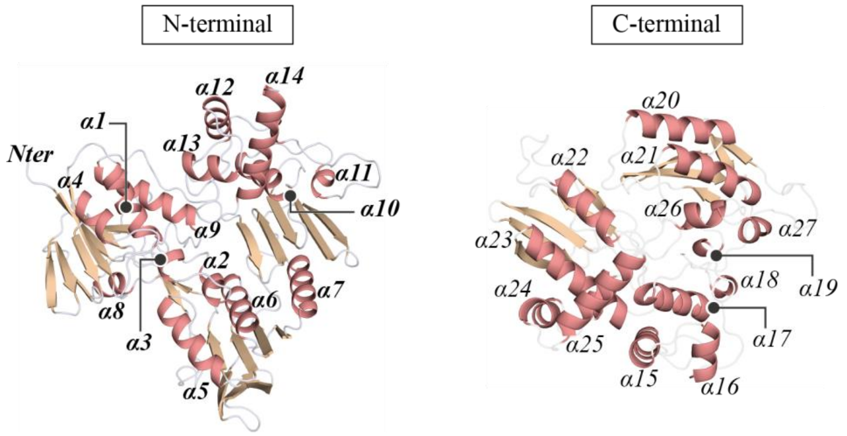


**Supplementary Fig. 7 Cartoon representation of the NADH-dependent nitrite reductase large subunit.** The N- and C-termini are located on the left and right, respectively. The α-helices numbered sequentially from the N- to C-terminus.

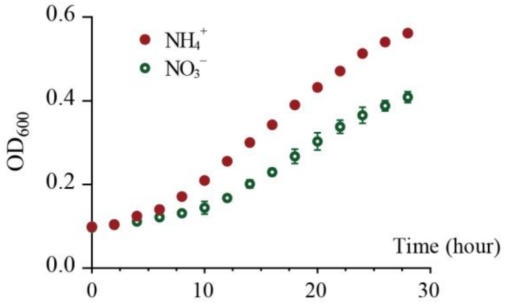


**Supplementary Fig. 8 Growth curves of *P.marina* NBRC 103166 under different culture conditions.** Red data points depict growth with ammonium as the nitrogen source under standard conditions; green data points depict growth under anaerobic conditions with nitrate as the sole nitrogen source.

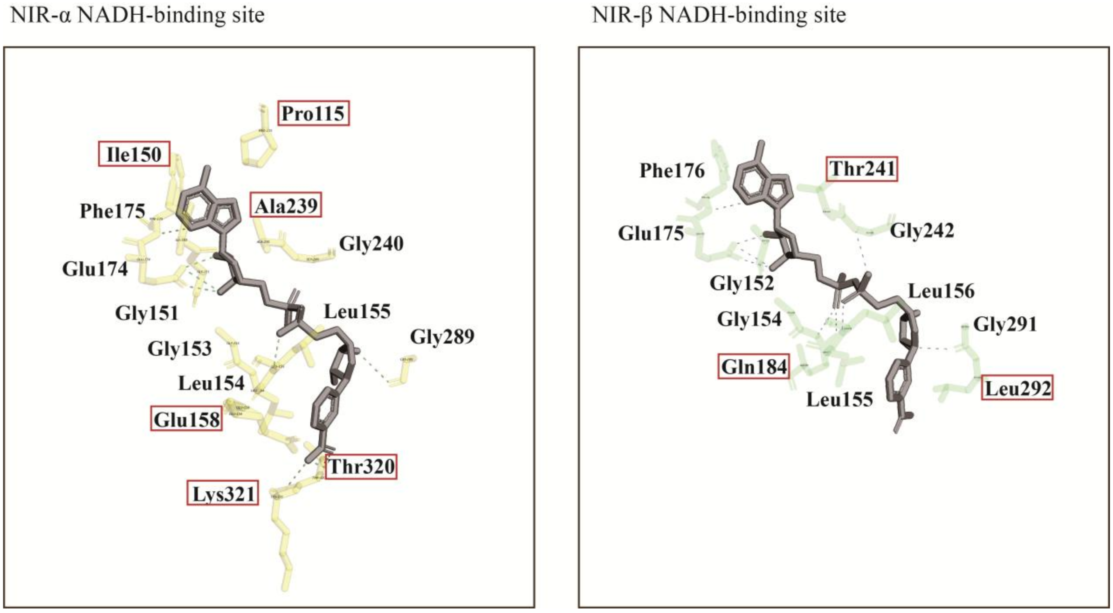


**Supplementary Fig. 9 Structural differences in the NADH binding sites between the two NirB isoforms.** The NADH molecule is shown in dark grey. Amino acid residues involved in NADH binding are depicted in stick representation: yellow for NIR-α (WP_028868406.1) and light green for NIR-β (WP_028868177.1). A red rectangle highlights the divergent residues between the two binding sites.

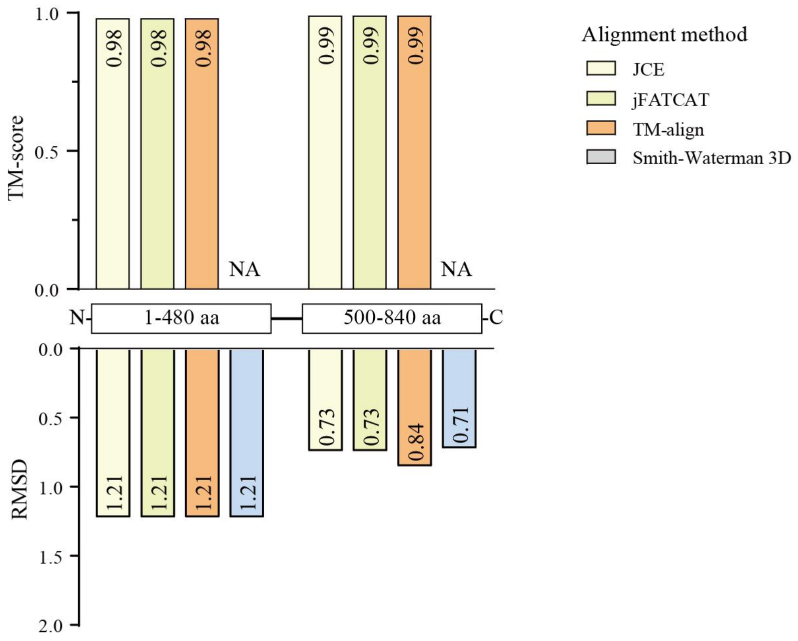


**Supplementary Fig. 10 Structural comparison of NADH-dependent nitrite reductase large-subunit models derived from AlphaFold2 and SeedFold.** To mitigate distortion from flexible regions, the N- and C-terminal domains were aligned and evaluated separately. To compare the structural models, the NADH-dependent nitrite reductase large-subunit structures predicted by AlphaFold2 and SeedFold were aligned using four independent methods: JCE, jFATCAT, TM-align, and Smith-Waterman 3D. The resulting alignments were evaluated by TM-score and RMSD. Our analysis revealed minimal structural differences between the models generated by the two prediction methods.
